# Molecular signatures of longevity identify compounds that extend mouse lifespan and healthspan

**DOI:** 10.1101/2025.06.26.661776

**Authors:** Anastasia V. Shindyapina, Alexander Tyshkovskiy, Perinur Bozaykut, José P. Castro, Maxim V. Gerashchenko, Alexandre Trapp, Margarita V. Meer, Bohan Zhang, Jesse R. Poganik, Steve Horvath, Richard A. Miller, Vadim N. Gladyshev

## Abstract

Longevity interventions in mammals are typically discovered on a case-by-case basis, hindering systematic geroprotector development. We developed a platform for the identification of longevity interventions integrating longevity gene expression biomarkers within and across species, in silico chemical screening, analyses of selected compounds in cell culture, short-term dietary interventions coupled with omics profiling, and ultimately lifespan studies in mice. This approach identified compounds (selumetinib, vorinostat, celastrol, AZD-8055, LY-294002) that extended lifespan and/or healthspan in aged C57BL/6JN male mice, with limited effects in females. In addition, selumetinib and vorinostat increased lifespan when administered to young, genetically heterogeneous UM-HET3 mice. Our biomarker-driven platform accelerates geroprotector discovery, offering a scalable approach to target conserved longevity pathways.

## Introduction

Targeting the aging process has the potential to postpone the onset of multiple age-related diseases simultaneously and to extend lifespan^1^. However, despite extensive research efforts in the past decade, there are no definitive interventions proven to extend human lifespan. To streamline the assessment of pharmacological interventions in human clinical trials targeting aging, it is useful to initially establish safety and efficacy profiles in preclinical models, where mouse is the most convenient model due its relatively short lifespan and lower housing cost compared to other mammals. Subsequently, interventions that demonstrate safety and efficacy in mice may subsequently be evaluated in human clinical trials.

Several interventions, including pharmacological, lifestyle and genetic, have consistently demonstrated the ability to extend the lifespan of mice. Rapamycin, for instance, extends the median lifespan of both male and female mice by up to 26% whether administered early or late in life^2,3,4,5^. Genetic manipulation of growth hormone signaling can extend mouse lifespan by up to 40%, across sexes and genetic backgrounds^6,7,8^. There is also evidence that reduced growth hormone signaling improves health outcomes in humans by lowering cancer incidence rates^9^. Another gold-standard intervention is calorie restriction (CR). Over a century ago CR was shown to extend rat lifespan^10^, and for many years it has been a well-established intervention in mice^11^. Additionally, CR was the first intervention shown to extend the healthspan of rhesus monkeys, providing the first evidence that the effect of life-extending intervention can be translated from rodents to primates^12^.

Despite the success of these and other longevity models, they have all been discovered on a case-by-case basis, hand-picked across many possible pharmacological, genetic, dietary and lifestyle interventions. Testing them on a larger scale remains a challenge due to cost and time involved. The cost of a well-powered study measuring mouse lifespan typically exceeds $100,000, mostly driven by the expense of mouse housing. Screening for anti-aging drugs in short-lived organisms (e.g. *C. elegans*, yeast and Drosophila) can be performed rapidly and at much lower cost. However, interventions identified by these screening studies may have more limited applicability to humans due to lower tissue complexity and a smaller number of conserved proteins targeted by these interventions compared to mice.

We previously identified biomarkers of longevity that include gene expression signatures of lifespan-extending interventions^13^ and long-lived mammalian species^14^. Using these signatures, we established a scalable platform for prediction of compounds with the potential to extend lifespan and healthspan. We have now carried out an *in silico* screen on more than 4,000 compounds, followed by *in vitro* testing of top 111 hits in primary human hepatocytes (PHH) and *in vivo* testing of 25 most promising candidates in mice. This approach enabled the identification of several compounds that improved health and extended lifespan when administered to young and/or old mice.

## Results

### Platform for the identification of longevity interventions based on molecular profiling

To predict compounds with potential lifespan-extending properties, we utilized gene expression-based longevity signatures that reflect molecular changes induced by established lifespan-extending interventions in mice^13^ and those characteristic of long-lived mammalian species^14^. We (i) conducted an *in silico* screen of over 4,000 compounds in the Connectivity Map (CMAP) database^15^, examining the association of their gene expression profiles in human cells with longevity biomarkers. We then (ii) selected 111 top candidates and profiled the transcriptomes of PHH treated with these molecules. We further (iii) selected 25 compounds for *in vivo* application, subjecting UM-HET3 male mice to diets containing each of these for 1 month and performing RNA-sequencing of the livers and kidneys of these mice. Finally, we tested (iv) the top 10 compounds associated with signatures of longevity across biological models for their effects on lifespan and healthspan in 24-25 month-old C57BL/6JN mice, and (v) 3 compounds in UM-HET3 mice by starting treatment at a young age and measuring lifespan, biomarkers of aging and multiple health parameters (Fig. 1A, Table S1). We also applied aging clocks to validate the effect of identified compounds on biomarkers of aging. Below, we describe this approach in detail.

**Figure 1.**
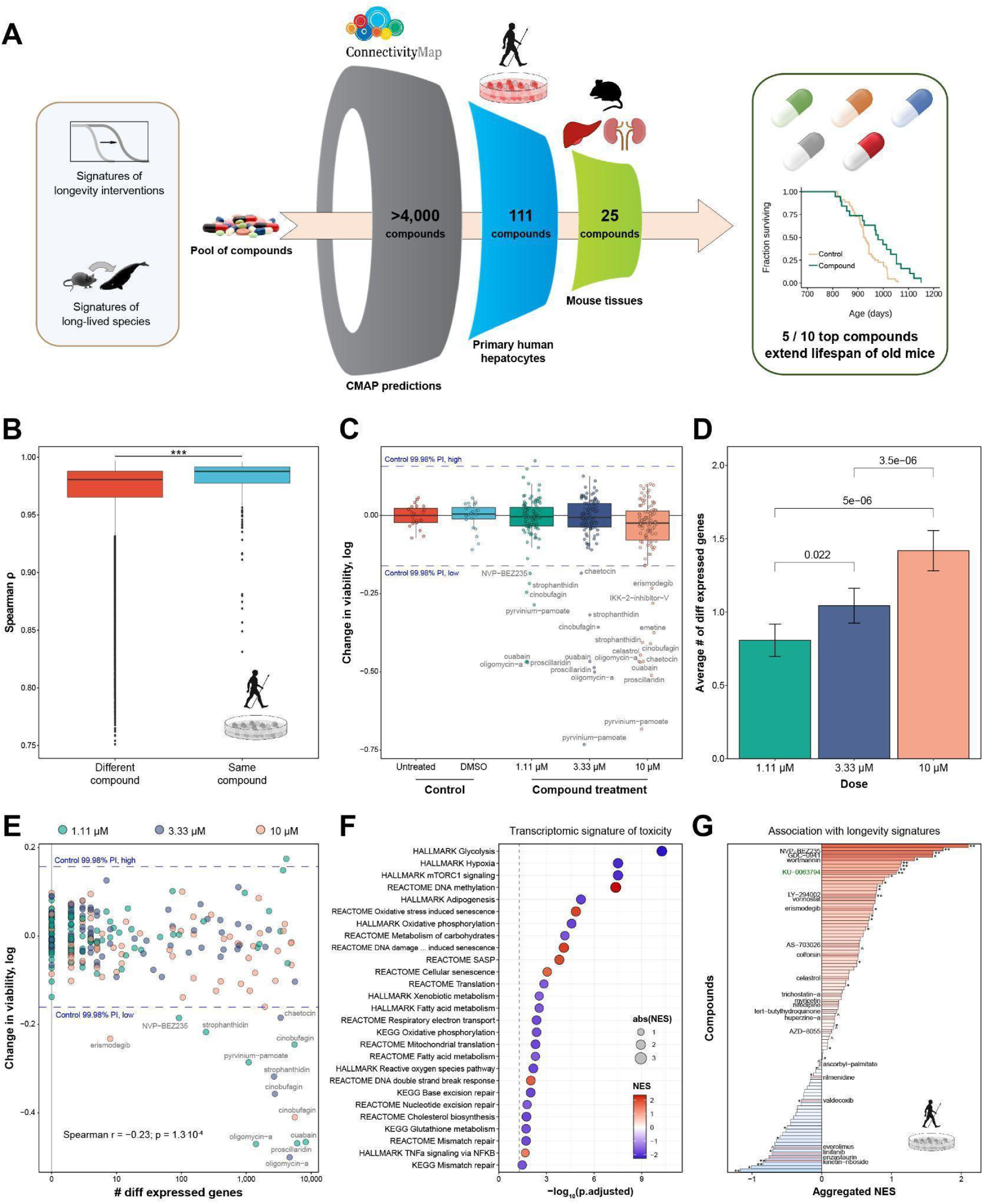
Screening of compounds with longevity-associated gene expression profiles in mammalian cell lines. **(A) Scheme of the screening platform.** Effect of compounds on gene expression was evaluated using developed biomarkers of lifespan-extending interventions and long-lived mammalian species. Tested biological models include human cell lines from Connectivity Map, primary human hepatocytes (PHH), and tissues from adult UM-HET3 male mice subjected to diets containing the identified compounds for 1 month. Compounds positively associated with longevity biomarkers across models were tested for the effect on survival in old mice. **(B) Pearson correlation between gene expression profiles of PHH subjected to different compounds (left) or different doses of the same compound (right).** Difference in median correlation between profiles of the same and different compounds was compared with Wilcoxon rank sum test. **(C) Effect of compounds on PHH viability.** Dots represent individual biological replicates untreated (red), treated with DMSO (lightblue) or treated with different concentrations of chemical compounds. 99.98% prediction interval for viability of control cells is shown with a blue dotted line. Compounds leading to decrease of viability below 99.98% prediction interval are labeled in gray. n=1 for each dose of compound, n=23 for DMSO-treated cells, n=24 for untreated cells. **(D) Average number of genes differentially expressed in PHH in response to various doses of chemical compounds compared to untreated and DMSO-treated cells.** Genes were considered differentially expressed if adjusted p-value < 0.05. n=3 for each dose of compound, n=67 for DMSO-treated cells, n=69 for untreated cells. Data are mean ± SE. Pairwise differences in average number of differentially expressed genes across doses were assessed with the Wilcoxon signed-rank test. BH-adjusted p-values are indicated with labels. **(E) Dependence between effect on PHH viability and number of differentially expressed genes.** Dots represent individual dose-compound groups and are colored based on compound concentration. 99.98% prediction interval for viability of control cells is shown with a blue dotted line. Compounds leading to decrease of viability below 99.98% prediction interval are labeled in gray. Spearman correlation coefficient between viability and number of differentially expressed genes and corresponding p-value are shown in text. **(F) Pathways enriched for genes associated with cell death (toxicity) in PHH assay.** Pathway enrichment analysis was performed with GSEA using HALLMARKS, KEGG, and REACTOME ontologies. BH adjusted p-value (in log scale) is shown on x axis, and normalized enrichment score (NES) is reflected with color and size of dots. **(G) Aggregated longevity association score of gene expression changes induced by chemical compounds in PHH.** Compounds are ranked by their average NES based on the signature of maximum lifespan in mice and signature of long-lived species across organs. Compounds selected for screening in mouse tissues after 1 month of treatment are labeled and indicated by red lines. Statistical significance of association (BH-adjusted harmonic mean p-value) is denoted by asterisks. KU-0063794, established to extend lifespan in old mice in our previous study, is labeled in green. The output of the association test is in Table S2. ^ p.adjusted < 0.1; * p.adjusted < 0.05; ** p.adjusted < 0.01; *** p.adjusted < 0.001. DMSO: Dimethyl sulfoxide; PHH: Primary Human Hepatocytes; CMAP: Connectivity Map; NES: Normalized Enrichment Score. See also Figure S1 and Tables S2, S3.

### Screening of compounds for their association with transcriptomic signatures of longevity in human cells

First, we screened for compounds that trigger gene expression changes similar to established longevity interventions in human cell lines using the CMAP database^15^. CMAP contains transcriptomic profiles of cell lines exposed to thousands of compounds, enabling the identification of perturbagens that produce gene expression changes aligned with the desired longevity-related signatures. As query inputs, we utilized four transcriptomic signatures of mammalian longevity derived from liver, kidney, or brain, and or pooled data across organs^14^, along with five transcriptomic signatures of established lifespan-extending interventions in mice, including CR, rapamycin, mutations associated with growth hormone (GH) deficiency, and biomarkers of maximum and median lifespan across interventions^13^. We ranked the perturbagens based on their aggregated longevity scores, defined as the mean of normalized enrichment scores (NES) estimated from signatures of maximum lifespan in mice and long-lived species across organs (Fig. S1A-B; Table S2A). In agreement with our previous studies, the association scores were generally positively correlated across signatures of long-lived species, and the majority of lifespan-extending intervention signatures produced consistent predictions (Fig. S1B-C).

To independently validate the gene expression profiles of compounds from CMAP, we selected 111 molecules, prioritizing those with high aggregated association scores (Fig. S1A), and tested their effects on gene expression in PHHs (Table S3). Each compound was dissolved in DMSO and applied to the PHHs at 3 concentrations: 1.11 µM, 3.33 µM and 10 µM. Untreated cells and PHHs exposed only to DMSO were used as controls. Gene expression profiles of hepatocytes treated with different concentrations of the same compound were more similar to each other than to the profiles of hepatocytes treated with different compounds (Fig. 1B), suggesting that these compounds produce distinct molecular footprints consistent across doses.

To evaluate potential compound-induced toxicity, we assessed the viability of PHHs in each condition using the CellTiter-Glo assay. As expected, the highest concentration (10 µM) resulted in significantly reduced median cell viability compared to control PHHs (Mann-Whitney U test p-value = 0.006) (Fig. 1C). Additionally, we observed a dose-dependent increase in the number of compounds that caused substantially lower viability compared to control cells (Bonferroni adjusted p-value < 0.01). Notably, some compounds, such as oligomycin-a, pyrvinium-pamoate, ouabain and cinobufagin, reduced cell viability across all tested concentrations.

In general, more genes were differentially expressed (BH-adjusted p-value < 0.05) in response to increased doses of test agents (Fig. 1D). The number of differentially expressed genes per compound was negatively correlated with cell viability (Spearman ⍴ = −0.23) (Fig. 1E), although considerable reorganization of the cellular transcriptome did not always result in a noticeable decline in viability. To characterize the molecular signatures of chemical-induced toxicity in primary hepatocytes, for each gene we evaluated the correlation of its expression and cell viability. Over 8,000 genes were significantly correlated with cell death (BH-adjusted p-value < 0.05) (Fig. S1D). Notably, the top pathways enriched for genes upregulated in response to toxic perturbations included DNA methylation, DNA damage and oxidative stress induced senescence, senescence-associated secretory phenotype (SASP), DNA double strand break response, and TNF-α signaling via NF-κB (Fig. 1F), highlighting the accumulation of molecular damage leading to cell death. In contrast, genes involved in energy metabolism, oxidative phosphorylation, translation, and mTORC1 signaling were downregulated, indicating an overall suppression of cellular biochemical and physiological processes under toxic conditions.

To select candidate compounds for subsequent assessment in mouse organs, we eliminated toxic treatments and molecules lacking common names. We then identified gene expression signatures for each remaining compound across different doses, and assessed their association with the transcriptomic biomarkers of longevity (Fig. S1E; Table S2B). Similar to the CMAP results, association scores were positively correlated within the sets of biomarkers derived from longevity interventions and long-lived species, but not between these groups (Fig. S1F). This finding suggests that individual treatments and long-term evolutionary processes may affect distinct molecular mechanisms of lifespan regulation, as indicated in our previous study^14^. To examine whether the effects of compounds on transcriptomic biomarkers of longevity are consistent between cell cultures and animal tissues, we selected 23 interventions with varying aggregated longevity scores (Fig. 1G), all of which had published experimental protocols for administration in mice via diet or oral gavage. Additionally, we included two molecules (selumetinib and pyrvinium-pamoate) that demonstrated positive associations with certain longevity signatures in CMAP and for which safe dosages for oral administration in mice had been established (Table S1, S2A). The resulting compounds were subsequently tested in young, genetically heterogeneous mice *in vivo*.

### Short-term treatment of mice with the selected compounds

We subjected 3-month-old UM-HET3 male mice to 25 selected compounds for 1 month via diet. All compounds had previously been tested in mice for other effects, allowing us to select a safe dosage for each (Table S1). Animals were weighed immediately before treatment, two weeks after treatment and at the end of the 1-month treatment period. Two compounds, AS-703026 and wortmannin, resulted in over 10% weight loss in the animals and were subsequently excluded from further testing. Wortmannin also caused heart enlargement in the treated animals.

To identify gene expression changes induced by the tested compounds in mouse tissues, we performed RNA-seq of the liver and kidney samples from treated and age-matched control males. Using an FDR threshold of 0.05, we identified differentially expressed genes for each compound in individual tissues as well as those combined (Fig. 2A). Of the 25 compounds tested, 23 induced significant differential expression of at least one gene in at least one of the analyzed settings - liver, kidney, or across tissues. Interestingly, AS-703026 and wortmannin, which were toxic at the selected doses used, produced the highest number of differentially expressed genes in the tested organs, especially in the liver, consistent with the dependence between toxicity and the effect size on transcriptome previously observed in PHHs (Fig. 1E).

**Figure 2.**
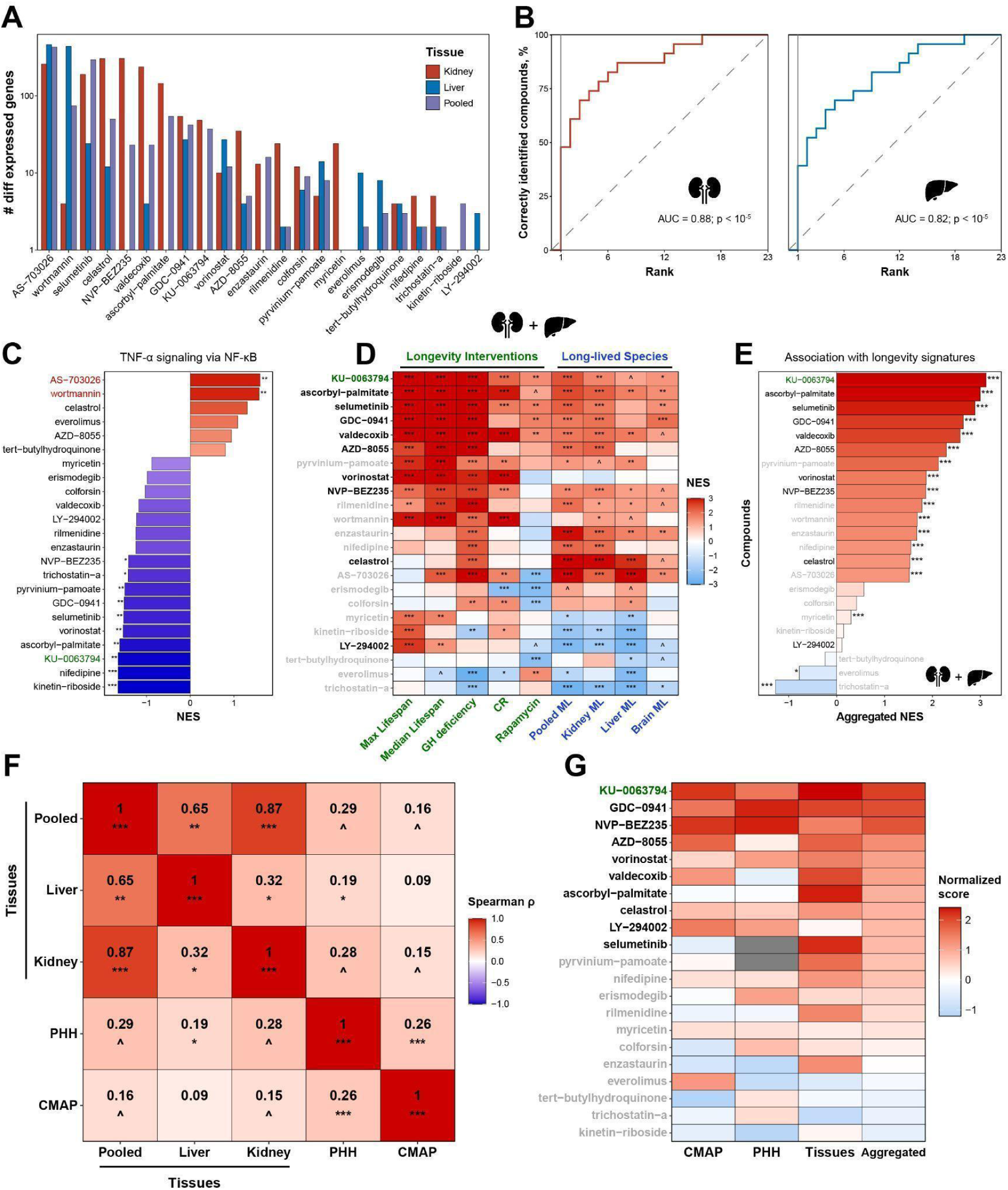
Screening for compounds with longevity-associated transcriptomic profiles in tissues of UM-HET3 male mice after short-term application. **(A) Number of differentially expressed genes perturbed by compounds after 1 month of treatment in murine liver (blue), kidney (red) and pooled across tissues (purple)** 23 out of 25 compounds induced differential expression of at least one gene in at least one tissue group after 1-month oral treatment. n=4 per tissue for treated mice; n=6 per tissue for control samples. **(B) Consistency between gene expression changes induced by individual compounds in the liver and kidney.** The y axis reflects the percentage of compounds, whose gene expression profiles in kidney (left) and liver (right) are among the top correlated with the reference profiles of the same molecules in the other organ. The x-axis indicates the number of top-ranked positions used to identify a match in the ranking list. Pearson correlation was used to rank the compounds. AUC and permutation test p-value are shown in text. **(C) Enrichment of gene expression changes induced by individual compounds across tissues for genes associated with TNF-α signaling via NF-κB.** Pathway enrichment analysis was performed with GSEA. Gene set was derived from HALLMARK ontology. Bars are colored based on normalized enrichment scores (NES). BH-adjusted p-values are denoted by asterisks. Toxic compounds and KU-0063794, established to extend lifespan in old mice in our previous study, are labeled in red and green, respectively. The whole list of enriched functions is in Table S3. **(D) Association of gene expression changes induced by tested compounds in murine tissues with signatures of lifespan-extending interventions (green) and long-lived mammalian species (blue).** Cells are colored based on the NES calculated for gene expression profiles of compounds across organs (kidney and liver). Signatures of maximum lifespan in mice (“Max Lifespan”) and long-lived species across organs (“Pooled ML”) were used to calculate the aggregated longevity score (presented on Fig. 2E). Compounds chosen for a survival study in old C57BL/6JN mice are labeled in black. BH-adjusted p-values are denoted by asterisks. The output of the association test is in Table S2. **(E) Aggregated longevity association score of gene expression changes induced by compounds in murine tissues.** Compounds are ranked by their average NES. Compounds selected for a survival study in old C57BL/6JN mice are labeled in black. Statistical significance of association (BH-adjusted harmonic mean p-value) is denoted by asterisks. **(F) Aggregated Spearman correlation of longevity signature association scores induced by chemical compounds across various biological models.** Pairwise Spearman correlation coefficients were calculated separately for each longevity signature and aggregated using a random-effect model. Aggregated correlation coefficients and corresponding BH-adjusted p-values are shown with text and asterisks, respectively. **(G) Normalized aggregated longevity association scores of compounds tested in murine tissues and human cells.** Cells are colored based on scaled association scores calculated separately for each biological model, including human cell lines from CMAP, PHH, and murine tissues. Compounds are sorted by the weighted average longevity association score across all tested biological models (“Aggregated”). Compounds chosen for a survival study in old C57BL/6JN mice are labeled in black. Missing data is indicated in grey. ^ p.adjusted < 0.1; * p.adjusted < 0.05; ** p.adjusted < 0.01; *** p.adjusted < 0.001. AUC: Area Under the Curve; Max: Maximum; CR: Caloric restriction; GH: Growth hormone; ML: Maximum Lifespan; PHH: Primary Human Hepatocytes; CMAP: Connectivity Map. See also Figure S2 and Tables S2-S3.

To test whether compounds induce similar molecular phenotypes across different murine organs, we compared gene expression changes in the liver and kidney for each intervention. For each compound, we calculated the correlation between its gene expression profile in one tissue and the profiles of all compounds in the other tissue. We then ranked the treatments based on their correlation coefficients with the selected compound and identified the position of that same compound within the list. We found that half of the drugs ranked among the top two molecules with the highest correlation coefficients when comparing profiles across tissues (Fig. 2B). The area under the curve for such a simple classification algorithm was 0.88, and it was 0.82 when average changes induced by compounds in the kidney and liver were sorted based on the reference profile in the other tissue, respectively (permutation p-value < 10^-5^). This indicates the existence of a shared effect of interventions on transcriptomes in different tissues, which may be related to similar molecular mechanisms of action for each drug across cell types or to shared environmental effects that influence different tissues similarly via extracellular communication.

Surprisingly, although hepatocytes represent the predominant cell type in the liver, we did not observe significant consistency between gene expression changes induced by the same compound in PHHs and mouse liver (Fig. S2A) or kidney (Fig. S2B). This finding suggests that the transcriptomic response of the tissue to a particular compound may not directly reflect the intervention’s effect but rather the impact of the tissue environment systemically perturbed by this intervention. Another possibility is that the drugs require conversion into their active forms by enzymes present in animals but absent in cultured cells. These hypotheses may explain why different tissues within the same organism demonstrate similar gene expression responses to compounds, while nominally similar cell types, e.g. hepatic cells, respond differently when subjected to the compound directly or administered systemically.

To characterize cellular pathways associated with toxicity at the gene expression level in murine tissues, we performed functional enrichment analysis of the transcriptomic changes induced by each compound across organs, and searched for enriched pathways that distinguish the effects of toxic compounds (AS-703026 and wortmannin) from those of others (Fig. S2C). The top term significantly upregulated in response to these interventions (BH-adjusted p-value < 0.05) but not to other treatments was TNF-α signaling via NF-κB (Fig. 2C), pointing to the conserved association of this pathway with the accumulation of molecular damage in mammalian organs and cell lines. We also found several other pathways associated with toxicity both in mouse tissues and PHHs, such as cellular senescence and fatty acid metabolism, upregulated and downregulated in response to toxic compounds, respectively (Fig. S2C).

To evaluate the potential effects of non-toxic compounds on longevity biomarkers in murine tissues, we assessed the association of their differential expression profiles with the signatures of established lifespan-extending interventions and mammalian longevity (Fig. 2D-E). Similar to the PHH and CMAP models, the association scores estimated based on interventions and species longevity showed a positive correlation within each group (Fig. S2D). In addition, we observed a statistically significant, moderate positive correlation in association scores across organs (Fig. 2F; Fig. S2E-F), suggesting partial conservation of molecular profiles induced by chemical treatments across tissues, while also highlighting the presence of tissue-specific variation. To rank compounds by their association with longevity biomarkers in mouse organs, we calculated aggregated scores, averaging the signatures of maximum lifespan in mice and multi-tissue signatures of long-lived species (Fig. 2E). Interestingly, some compounds, such as KU-0063794, ascorbyl-palmitate, selumetinib, GDC-0941, valdecoxib and AZD-8055, induced gene expression changes associated with extended lifespan according to both biomarkers of longevity within and across species (Fig. 2D), pointing to the promising potential of these molecules for promoting longevity. Notably, most of these compounds also resulted in significant downregulation of TNF-α signaling (Fig. 2C). On the other hand, some compounds, such as LY-294002, exhibited more diverse effects, inducing changes positively associated with the signatures of established lifespan-extending interventions, especially in the liver, but negatively associated with the signatures of long-lived mammalian species, especially in the kidney (Fig. 2D; Fig. S2E-F).

We then examined the consistency of compound association scores derived from gene expression data across various biological models (Fig. 2F). Interestingly, although we did not observe significant similarity in the overall transcriptomic changes induced by the same compounds in primary human hepatocytes and mouse livers (Fig. S2A), their associations with signatures of longevity were positively correlated (Fig. 2F). On the other hand, CMAP predictions showed a statistically significant positive correlation only with the association scores in PHHs, while their correlation with scores of corresponding compounds in murine tissues were much weaker, suggesting that the effect of treatment on molecular profiles of immortalized human cells and mouse tissues may be too distinct, even when only the longevity-associated component is taken into account.

Finally, for the 21 compounds that induced differential expression of at least one gene without pronounced toxicity in mice after one month of treatment, we computed a meta-score by aggregating longevity association scores across all three biological models (CMAP, PHH, and murine tissues). We then sorted compounds prioritizing those supported by multiple models to reduce the likelihood of false positives (Fig. 2G). Notably, several compounds, such as KU-0063794, GDC-0941, NVP-BEZ235, vorinostat, and celastrol demonstrated positive longevity scores across all 3 models, and KU-0063794, the compound with the highest meta-score, had already been shown to significantly extend lifespan and frailty of old male mice in our earlier study^14^. To further validate the predictive power of our screening platform, we selected nine compounds with the highest meta-scores for testing, including PI3K inhibitors GDC-0941, NVP-BEZ235, and LY-294002, an mTOR inhibitor AZD-8055, a histone deacetylase (HDAC) inhibitor vorinostat, a COX-2 inhibitor valdecoxib, an antioxidant ascorbyl-palmitate, an anti-inflammatory natural compound celastrol, and a MEK1/2 inhibitor selumetinib. We then tested their effects on lifespan and healthspan of aged C57BL/6JN mice.

### Five out of ten predicted compounds extend lifespan or healthspan of old C57BL/6JN mice

To evaluate longevity properties of these compounds *in vivo*, we subjected twenty to twenty-four 25-month-old C57BL/6JN mice (∼10 males and ∼10 females) to each of 9 compounds, using 44 untreated animals (21 males and 23 females) as the control group. Prior to treatment, we recorded mouse body weight and their frailty index (FI) scores. We then assigned mouse cages to treatment groups such that the mean and deviation of weight and FI were similar across groups (Fig. S3A,E). Each group was then subjected to diets containing the same concentration of compounds as that tested in young UM-HET3 mice (Table S1). In addition to KU-0063794 validated previously^14^, four compounds significantly extended median lifespan: LY294002, AZD8055, celastrol and selumetinib (Fig. 3A-D); the other five compounds had no effect on lifespan (Fig. S3B, Table 1). LY294002 and AZD8055 showed a sex-specific effect, extending lifespan only in males. Celastrol and Selumetinib showed a trend toward extending lifespan in both males and females, although the study was under-powered to detect significance for each sex separately. None of the tested compounds had a significant impact on the animals’ body weight (Fig. S3A,C).

**Figure 3.**
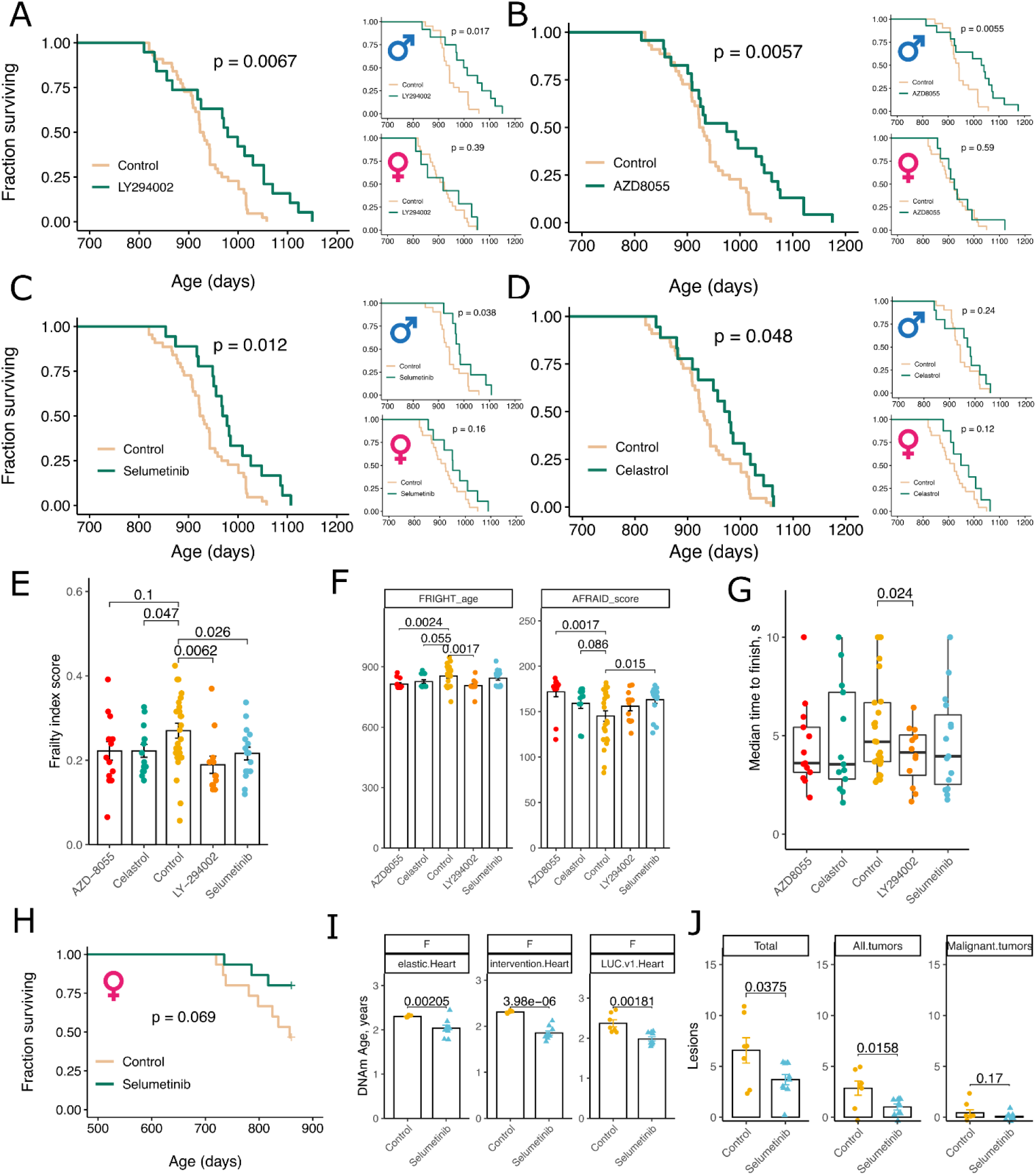
Lifespan and healthspan of aged C57BL/6JN mice treated by predicted longevity interventions. **(A)-(D) Lifespan of 24-25-month-old mice subjected to diets containing candidate longevity interventions.** Survival of both sexes, as well as males and females separately, when given diets containing 600 ppm LY294002, 20 ppm AZD8055, 100 ppm selumetinib or 8 ppm celastrol, all compared to untreated control. P-values were calculated with a log-rank test. **(E) Frailty index score measured in 31-month-old** C57BL/6JN **mice following 5 months on diets.** Mice were subjected to diets from 24-25 months of age and the frailty score was measured at 31 months of age in both sexes. P-values were calculated with a two-sided Student T-test. **(F) Frailty index clocks measured in 31-month-old** C57BL/6JN **mice after 5 months of treatment.** FRIGHT age predicts age (in days) of the mice based on frailty index measures, and AFRAID score predicts remaining lifespan, also in days. P-values were calculated with a two-sided Student T-test. Both males and females are included in the analysis. **(G) Median time to finish the gait speed test measured in 31-month-old** C57BL/6JN **mice after 5 months of treatment.** Gait speed was measured four times per mouse, and the median value is shown on the plot. P-values were calculated with a two-sided Student T-test. Both males and females are included in the analysis. **(H) Survival of female** C57BL/6JN **mice treated with 100 ppm selumetinib from 21 months of age.** P-values were calculated with a log-rank test. **(I) DNA methylation age of the heart measured in 27-month-old female** C57BL/6JN **mice given selumetinib.** Estimation of DNA methylation age in hearts of mice using three types of epigenetic clocks for heart tissue. P-values were calculated with a two-sided Student T-test. **(J) Number of lesions in cross-sectional necropsy study of 27-month-old female** C57BL/6JN **mice treated with selumetinib.** Total number of lesions, number of tissues with tumors and number of malignant tumors as diagnosed by pathologies from H&E staining of 11 organs during necropsy analysis. P-values were calculated with a two-sided Student T-test.

We further evaluated whether these compounds improved health by conducting non-invasive behavioral and frailty tests. The choice of tests and parameters was largely inspired by Bellantuono et al^16^. Each of the drugs that extended lifespan also reduced frailty index score, and improved age or lifespan predictors estimated by frailty index clocks (Fig. 3E-F). Furthermore, vorinostat improved frailty index scores and frailty clocks in males and females, despite lacking effect on lifespan of old C57BL/6JN mice (Fig. S3E). We also assessed gait speed as a measure of fitness. We previously showed that mouse gait speed decreases with age and correlates with rotarod and treadmill measures of muscle function^4^. LY294002, which extended lifespan, also improved gait speed in aged mice (Fig. 3G), and valdecoxib improved gait speed despite lacking life-extending benefits (Fig. S3F). Positive effects of LY294002 on mobility fitness are consistent with recent findings that hyperactive mTOR in aged muscle is detrimental to muscle performance, which can be rescued by rapamycin, also an mTOR inhibitor^17,18^.

To further test the longevity benefits of the drugs, we applied epigenetic clocks, based on the Horvath Mammalian Methylation Array platform ^19^, to the liver, kidney and heart of mice subjected to selumetinib. We established a separate cohort of 21-month-old C57BL/6JN female mice and subjected them to a diet containing selumetinib and analyzed tissues six months later. We found evidence suggesting that selumetinib may have improved survival in this cohort (Fig. 3H), replicating the findings in 25-month-old C57BL/6JN mice (Fig. 3A). Selumetinib significantly reduced the epigenetic age of the heart as based on three different heart-specific clocks (Fig. 3I), but had no effect on the predicted age of the kidney and liver, suggesting that this compound might affect the aging DNA methylome in a tissue-specific way.

Finally, we analyzed causes of natural death of animals subjected to candidate longevity interventions. There was no significant difference in the proportion of lesions between treated and control groups measured at the end of their life (Table S4). Interestingly, pathology analyses in a cross-sectional cohort of selumetinib-treated animals revealed a lower burden of lesions, and, in particular, of tumors (Fig. 3J). Thus, the lifespan-extending effects of selumetinib are potentially explained by delaying development of age-related diseases rather than changing their spectrum in aged mice.

### Two out of three tested compounds extend lifespan and healthspan in male genetically heterogeneous mice when treatment is initiated at a young age

We examined three of the selected compounds for possible longevity effects in genetically diverse mice. For this experiment, we chose selumetinib, which extended lifespan of old mice in two independent cohorts, vorinostat, which failed to extend lifespan but improved healthspan in old males and females, and GDC-0941, which showed no effect on lifespan or healthspan in old mice. We assigned slightly over one hundred UM-HET3 animals per diet starting at 6 months of age and followed the animals until death. Both selumetinib and vorinostat extended the lifespan of male UM-HET3 mice (Fig. 4A-D, Table 2), while GDC-0941 had no longevity benefits (Fig. 4E-F, Table 2). Again, we observed sex-specific effects on lifespan, wherein male mice benefited from both treatments but females did not (Fig. 4A-D). Interestingly, vorinostat extended the lifespan of UM-HET3 mice, despite having had no effect in old inbred (i.e. C57BL/6JN) mice. Vorinostat, but not the other two drugs, also led to weight loss of both sexes (Fig. 4G).

**Figure 4.**
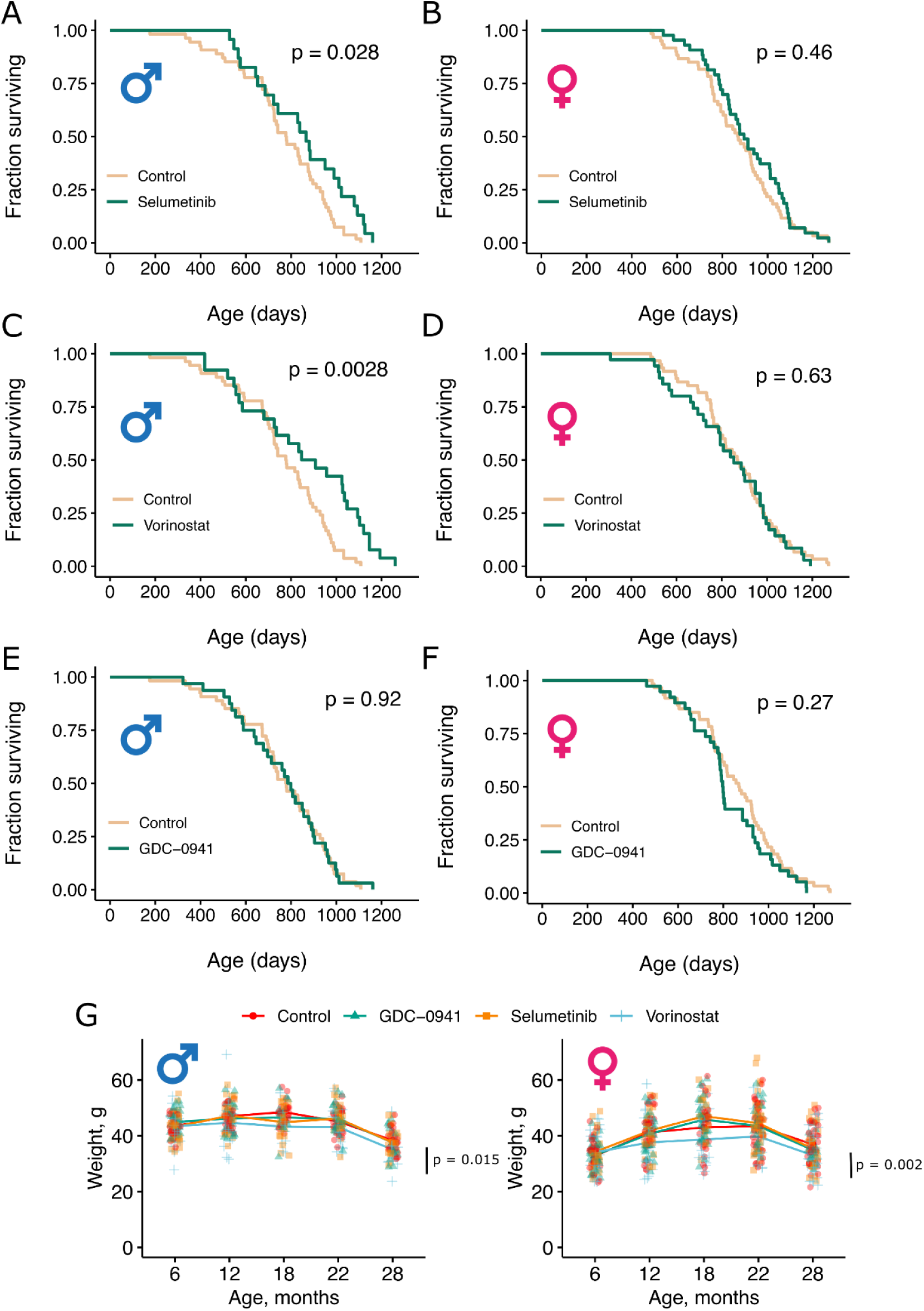
Lifespan of UM-HET3 mice treated by predicted longevity interventions starting at 6 months and across the whole lifespan. **(A)-(F) Lifespan of UM-HET3 male and female mice were subjected to diets containing 100 ppm selumetinib, 1000 ppm vorinostat, or 50 ppm GDC-0941** Treatment started at 6 months of age. P-values were calculated with a log-rank test. **(G) Body weight of UM-HET3 male and female mice treated with longevity compounds.**

After 9 months of treatment, we collected liver tissue from a subset of animals and performed RNA-seq to measure the long-term effects of compounds on the molecular profiles. Selumetinib and vorinostat induced differential expression of over 100 genes (adjusted p-value < 0.05) in each sex as well as across sexes (Fig. 5A). On the other hand, GDC-0941 treatment did not substantially affect the liver transcriptome, particularly in males, where only seven genes showed differential expression. In agreement with these findings, gene expression profiles of most GDC-041 samples clustered closely with those of the control group (Fig. 5B).

**Figure 5.**
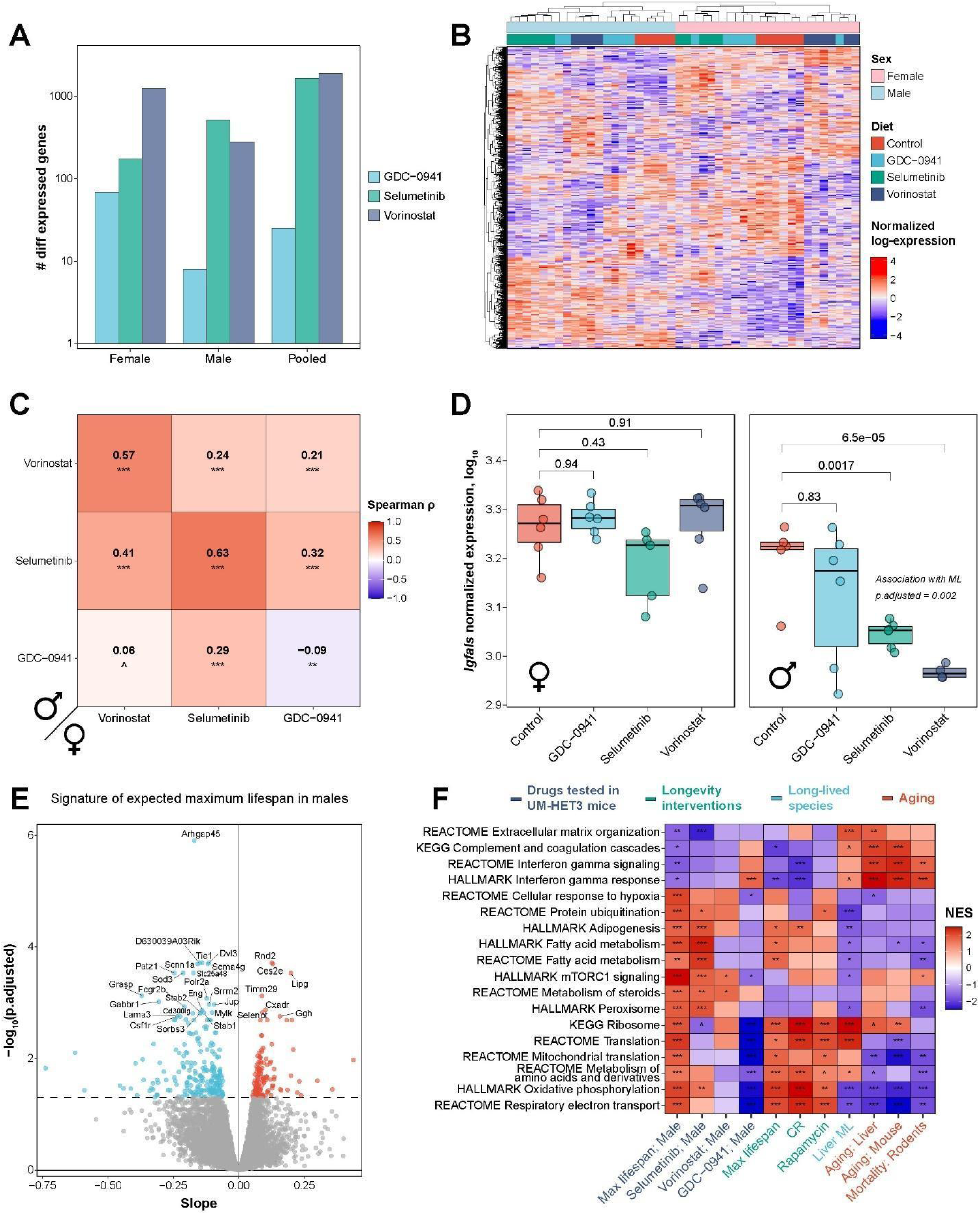
Liver gene expression profiles of UM-HET3 mice subjected to long-term treatment with the selected compounds. **(A) Number of differentially expressed genes perturbed by compounds in livers of 15-month-old mice following 9-month treatment** All 3 compounds induced differential expression of at least one gene in each sex compared to control mice. n=4-6 per tissue per sex. **(B) Heatmap of genes differentially expressed between the treatment groups.** Genes differentially expressed in at least one group in males or females according to ANOVA (adjusted p-value < 0.05) are presented. The x axis represents individual samples from a filtered dataset. Hierarchical clustering was performed for rows and columns using complete linkage and spearman correlation distance. Sex and treatment group of every sample is specified in color on the top. **(C) Spearman correlation of gene expression changes induced by compounds in livers of male (in rows) and female (in columns) mice from UM-HET3 strain.** For every pairwise comparison, the union of top 500 genes with the lowest p-value was used to estimate a correlation coefficient. Pairwise Spearman’s ⍴ are shown in color and text, and BH-adjusted p-values are denoted with asterisks. **(D) Normalized expression of *Igfals* in livers of age-matched female (left) and male (right) mice subjected to 9 months of oral treatment with selected compounds.** BH-adjusted p-values for pairwise comparisons between control and treated groups assessed with edgeR are labeled with numbers. Statistical significance of association between expression of the gene and maximum lifespan of male groups assessed with linear model is provided in text. **(E) Volcano plot of genes associated with maximum lifespan in male UM-HET3 mice across all experimental groups.** Top 30 genes with the lowest BH-adjusted p-value are labeled with gene symbols. FDR threshold of 0.05 is shown with dotted line, significant genes (BH-adjusted p-value < 0.05) positively and negatively associated with maximum lifespan are colored in red and blue, respectively. **(F) Functional enrichment of gene expression changes associated with expected maximum lifespan in UM-HET3 male mice across all treatment groups and with individual tested compounds (indigo), along with established signatures of lifespan-extending interventions (green), long-lived species (lightblue), aging and mortality (red).** GSEA was performed using HALLMARKS, REACTOME, and KEGG ontologies. Only pathways statistically significantly enriched for a signature of maximum lifespan in UM-HET3 male mice (BH-adjusted p-value < 0.05) are shown. Normalized enrichment score (NES) is shown with color, and BH-adjusted p-values are denoted by asterisks. The whole list of enriched functions is in Table S3. Max: Maximum; CR: Caloric restriction; GH: growth hormone; ML: Maximum Lifespan. ^ p.adjusted < 0.1; * p.adjusted < 0.05; ** p.adjusted < 0.01; *** p.adjusted < 0.001. See also Figure S4 and Table S3.

Principal component analysis (PCA) revealed two outlier samples with distinct transcriptomic profiles (Fig. S4A), which were excluded from further analysis. The remaining samples exhibited a clear separation based on the animals’ sex, indicating that sex-specific gene expression signatures were more pronounced than the effects of long-term treatment with these compounds. The transcriptomic changes induced by vorinostat and selumetinib showed a strong positive correlation between sexes (Fig. 5C), indicating a similar effect on the molecular phenotype in males and females. In contrast, we did not observe consistent signatures of GDC-0941 across sexes, likely due to its overall low impact on gene expression, particularly in males. Therefore, in general, vorinostat and selumetinib seem to induce similar effects on the mouse transcriptome, while the effect of GDC-0941 is too weak to clearly distinguish treated animals from controls. This may explain why this compound failed to produce a pronounced effect on health and lifespan.

To characterize conserved transcriptomic changes associated with drug-induced lifespan extension in males, we searched for genes whose expression levels correlated with expected maximum lifespan (90^th^ percentile) across all samples from the four treatment groups in UM-HET3 male mice (control, GDC-0941, selumetinib, and vorinostat). Utilizing a linear regression model and FDR threshold of 0.05, we identified 371 genes significantly associated with expected maximum lifespan across treatment conditions (Fig. 5D-E, Fig. S4B). Notably, they included several established biomarkers of mammalian lifespan^14^, such as *Igf1*, encoding insulin-like growth factor 1 (Fig. S4B), and *Igfals*, encoding its binding protein (Fig. 5D) (BH-adjusted p-value = 0.03 and 0.002, respectively). These associations further underscore the role of insulin signaling in the regulation of longevity.

To identify specific pathways associated with longevity in this mouse model, as well as the functions perturbed by individual treatments, we performed functional gene set enrichment analysis (GSEA) (Fig. 5F, Fig. S4C, Table S3). Among the top pathways enriched for genes associated with maximum lifespan in male UM-HET3 mice, we observed many established signatures of lifespan regulation in mice, including the upregulation of oxidative phosphorylation, fatty acid metabolism, cellular respiration, and mitochondrial translation. Conversely, genes involved in the complement and coagulation cascades, as well as interferon gamma signaling, were downregulated. Notably, most of these pathways are also affected by established lifespan-extending interventions, such as caloric restriction and rapamycin, while mouse aging and mortality are accompanied by the opposite gene expression changes.

Interestingly, several compounds demonstrated pronounced sex-specific effects at the level of aforementioned pathways. Thus, selumetinib significantly upregulated genes involved in oxidative phosphorylation in UM-HET3 males, while it had the opposite effect in females (Fig. S4C). However, some of its effects, including upregulation of genes associated with fatty acid metabolism and mTORC1 signaling, were consistent across sexes, suggesting that there may be a mixture of shared and sex-specific effects produced by the same compound in males and females. Notably, in addition to sex differences, compounds may also exert varying effects depending on the age of the animals and the duration of treatment. Thus, selumetinib and GDC-0941 induced similar gene expression changes in the livers of 15-month-old male mice after long-term treatment and in 4-month-old mice after short-term treatment (Fig. S4D). In contrast, vorinostat produced distinct gene expression profiles under these same conditions. This variability may help explain the differing effects on lifespan and healthspan observed for the same compounds across different mouse strains and experimental designs.

Alongside the survival study, we subjected 10-15 animals per group per sex to the same three diets, aging them and assessing their health (Fig. 6A). Frailty index scores were recorded in both lifespan and healthspan cohorts. Selumetinib reduced frailty index score in both males and females at 22 and 28 months (Fig. 6B,C) compared to control untreated groups. Conversely, an open field locomotor test revealed that vorinostat decreased physical activity in male mice that is characteristic of older mice, but not in females (Fig. 6D) in the 28-month age group, while GDC-0941 improved physical activity in 28-month-old female mice.

**Figure 6.**
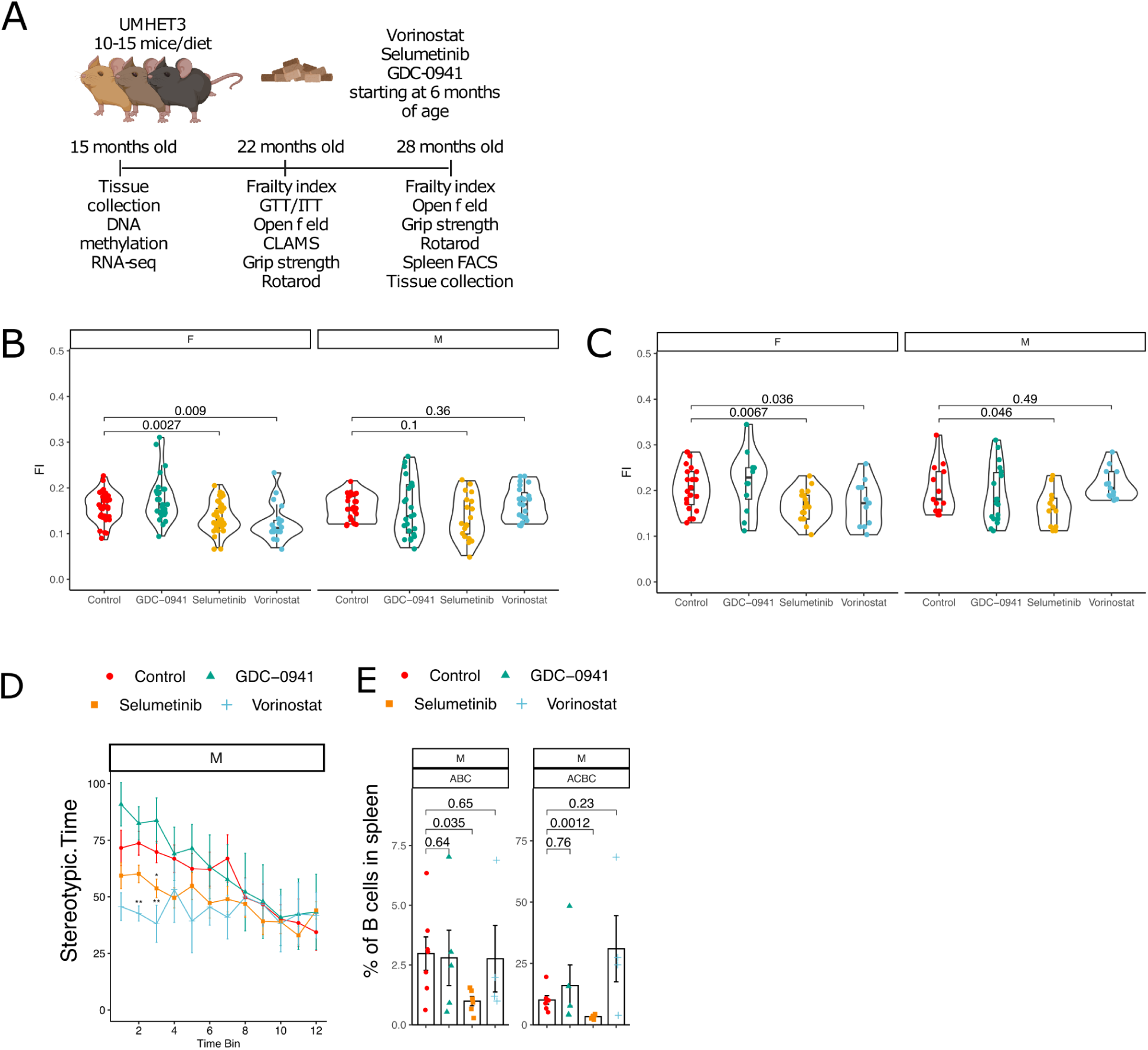
Measures of health in 22- and 28-months old UM-HET3 mice treated with longevity compounds starting at 6 months of age. **(A) Scheme of experiment to profile health in mice treated with longevity compounds.** **(B) Frailty index score measured in 22 and (C) 28-months old UM-HET3 mice** Females (F) and males (M) were treated with one of three longevity compounds and compared to untreated controls. P-values for only statistically significant results are shown (color-coded according to the diet), and were calculated with repeated measures ANOVA test. Each dot is a mouse, dots are colored and shaped based on the treatment group. **(D) Locomotor activity of old UM-HET3 mice treated with longevity compounds and compared to untreated controls.** Mean values with standard deviation are shown for each treatment group in males and females separately. Each bin is a 5-minute interval of 1-hour recording session. Stereotypic time is the time mice spend crossing the arena during recordings. **(E) Proportion of age-associated B cells (ABC) and age-associated clonal B cells (ACBC) measured in spleens of 28-months old UM-HET3 mice.** Mice were treated with one of three longevity compounds and compared to untreated controls. Each dot is a mouse, and dots are colored and shaped based on the treatment group. P-values were calculated with a two-sided Student T-test. ABCs were gated on live cells (Dapi-) as CD45+CD19+CD21-CD23-CD29+CD24-, and ACBC were gated as CD45+CD19+CD21-CD23-CD29+CD24+. **(G) Predicted epigenetic age of livers of 15-month-old UM-HET3 mice.** Each dot is a mouse, dots are colored based on the treatment group. P-values were calculated with a two-sided Student T-test.

Given that some longevity interventions affect the immune system, we measured the proportion of T cells, B cells and myeloid cells following treatment and found that they remained unchanged by any of the diets (Fig. S6A). Selumetinib substantially decreased the proportion of age-associated B cells and clonal B cells in male mouse spleens (Fig. 6E), which we previously showed to be associated with shorter lifespan and contribute to age-related lymphoma in mice^20^.

Finally, we tested the effect of drugs on metabolic health of the animals. Selumetinib improved glucose (Fig. 7A,B) and insulin (Fig. 7C,D) tolerance in 25-month-old UM-HET3 mice, regardless of sex. This effect was age-dependent since young mice treated with selumetinib were indistinguishable from the control group in the glucose tolerance test (GTT) (Fig. S5A,B). On the contrary, vorinostat induced glucose intolerance in an age-independent manner (Fig. S5A,B, Fig. 7A,B) in both sexes, and improved insulin tolerance only in old females (Fig. 7C,D). Metabolic cage records over 72 hours revealed that vorinostat and selumetinib had a profound effect on the respiratory exchange ratio (RER) of old males, but, similarly to the GTT, in the opposite direction (Fig. 7E,F). Selumetinib increased RER over controls, thus favoring carbohydrates as an energy source (Fig. S5E-F), while vorinostat decreased RER pointing to the use of fat as the main fuel source, which may explain weight loss in these animals. Strikingly, in both cases only male RER was affected by selumetinib and vorinostat (Fig. S5G-H). Interestingly, all three diets reduced activity of males (Fig. 7G-H) and females (Fig. S5E-F) at night, except for females that were treated with vorinostat. Both male and female mice on the vorinostat diet consumed less food (Fig. 7I-J, Fig. S5G-H), potentially explaining their weight loss.

**Figure 7.**
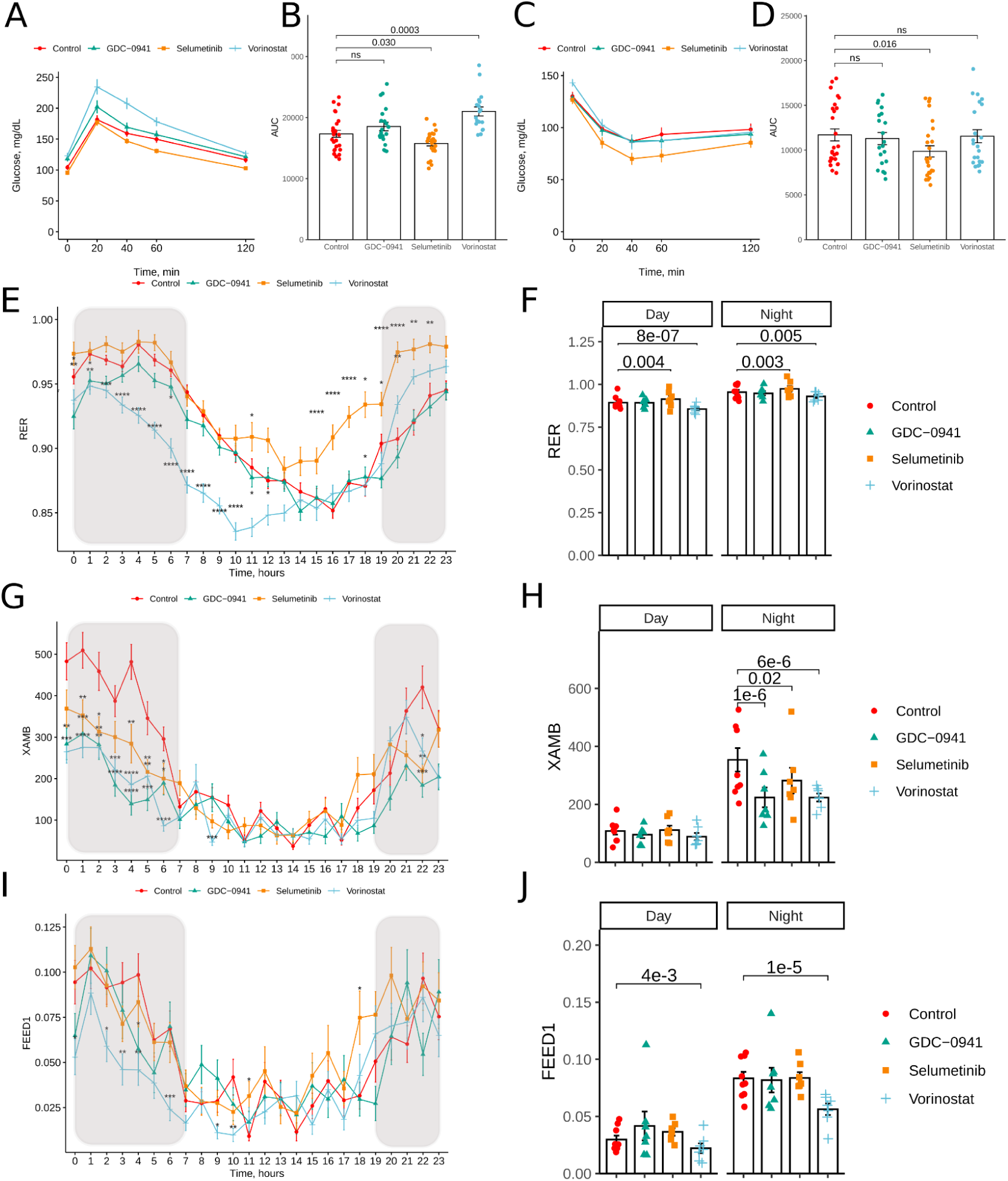
Measures of metabolic health in 22-25-months old UM-HET3 mice receiving treatments from 6 months of age. **(A-B) Results of glucose tolerance test measured in 22-months old UM-HET3 mice** Values are mean ± SD for both sexes combined. Area under the curve (AUC) for the glucose tolerance test. P-values are ANCOVA adjusted for sex. Each dot is a mouse. ns - not significant. **(C-D) Results of an insulin tolerance test measured in 22-months old UM-HET3 mice.** Values are mean ± SD for both sexes combined. Area under the curve (AUC) for the insulin tolerance test. P-values are ANCOVA adjusted for sex. Each dot is a mouse. ns - not significant. **(E-F) Respiratory exchange ratio (RER) of 25-months old UM-HET3 male mice treated with longevity intervention and compared to untreated control.** Values are mean ± SD, averaged over a 24-72 hour period of recordings. Shadowed areas indicate dark time at the facility, when mice are active. Barplots are averaged RER values during day and night periods. P-values are calculated with repeated measures ANOVA. Each dot is a mouse. **(G-H) Horizontal ambulatory activity (XAMB) of 25-months old UM-HET3 male mice treated with longevity intervention and compared to untreated control.** Values are mean ± SD, averaged over a 48-hours of recordings, after the first 24 hours when animals were adjusting to CLAMS. Shadowed areas indicate dark time at the facility, when mice are active. Barplots are averaged XAMB values during day and night periods. P-values on barplot are calculated with repeated measures ANOVA, and each dot is a mouse. **(I-J) Eating activity in grams (FEED1) of 25-months old UM-HET3 male mice treated with longevity intervention and compared to untreated control.** Values are mean ± SD, averaged over a 48-hours of recordings, after the first 24 hours when animals were adjusting to CLAMS. Shadowed areas indicate dark time at the facility, when mice are active. Barplots are averaged FEED1 values during day and night periods. P-values on barplot are calculated with repeated measures ANOVA, and each dot is a mouse.

Overall, selumetinib improved multiple health metrics in aged mice, including frailty index, grip strength, glucose and insulin tolerance, RER, and the burden of age-associated B cells. Vorinostat had limited effects on health parameters of mice with the exception of improved performance on rotarod, while some effects seemed detrimental (glucose intolerance and weight loss). GDC-0941 showed minimal effects on the overall health of animals, with the exception of improved rotarod performance and higher exploratory activity in females, yet lower physical activity overnight.

## Discussion

Here we show that transcriptomic responses to pharmaceutical compounds in young mice can predict their potential to extend lifespan and healthspan. Using the CMAP database we first selected compounds that elicit gene expression profiles similar to those associated with established longevity interventions and species longevity. This step allowed us to focus on hundreds rather than thousands of compounds. We tested 111 compounds in primary human hepatocytes and 25 molecules in young UM-HET3 mice to document short-term effects of treatment.

We found that our platform predicted lifespan-extending effects of compounds in aged C57BL/6 JN mice at 50% success rate (5 out of 10 treatments tested). Additionally, one more compound, vorinostat, improved the health of old C57BL/6JN mice and extended the lifespan of UM-HET3 mice when given earlier in life. Five of the lifespan-extending compounds identified in our assay had not been previously tested for longevity effects in mice. Some of these molecules target pathways that already have been shown to affect longevity in mice (e.g. AZD8055, an mTOR inhibitor, and selumetinib, a MEK1/2 inhibitor), mimicking other established lifespan-extending interventions, such as rapamycin^2,3,4,5^, another mTOR inhibitor, which has been shown to extend mouse lifespan alone and in combination with trametinib^21^. Other compounds have less characterized mechanisms (e.g., vorinostat, an HDAC inhibitor; LY294002, a PI3K inhibitor; and celastrol with multiple undefined targets). It is not unusual, however, for compounds to target multiple pathways, so selectivity of these compounds should be considered with caution.

While we observed an unusually high success rate in the current study, a subsequent study testing additional predicted compounds and combinations, currently in progress, is clearly not as effective. As such, conclusions on the success rate would be premature. Further studies are needed to assess how efficient platforms like ours are in predicting longevity interventions. It is clear, however, that they can predict such interventions.

An additional interesting observation is that the effects of compounds on gene expression varied between *in vivo* (tissue) and *in vitro* (cell culture) models. For example, we found that upon treatment, PHH gene expression changes did not show significant similarity with those in mouse tissues (liver and kidney). Nevertheless, we found that the PHH association with the signatures of longevity was positively correlated with those in mouse liver. At the same time, CMAP predictions showed a significant positive correlation solely with PHH, strongly suggesting that the compound elicited different transcriptome changes (in scope and in character) on human cells *in vitro* and mouse tissues *in vivo*. Additionally, we observed that the compounds more strongly altered gene expression in tissues than in cultured cells. Thus, most of the compounds failed to change gene expression profiles of primary hepatocytes at any of the three concentrations tested, while having profound effects on mouse liver. As such, our findings suggest that testing compounds in mice is more informative when screening for longevity compounds and should be prioritized over cell culture models, at least when gene expression is the primary readout. This might be due to several factors: (a) indirect effects may be at play via other tissues *in vivo*; and (b) there may be effects of conversion of a compound to an active agent, and/or excretion of the agent.

This issue becomes particularly important when screening is based on biomarkers identified at the tissue level, such as the molecular signatures of longevity utilized in this study. It might be most informative to test candidates primarily using short-term treatments of mice, if such tests are not cost-prohibitive and if the initial pool of compounds is limited. At the same time, liver and kidney largely overlap in their transcriptomic signatures in response to interventions. As compounds elicit similar responses in kidney and liver, the data suggest a possibility of using only one tissue of these two for lifespan extension potential, which would reduce the cost of future analyses, although predictions might be improved by adding tissues that contain more of proliferative cell types or performing very different functions (like adipose tissue or brain). While our screen focused on drugs and other chemicals, conceptually, the findings should apply for identifying other types of longevity interventions, e.g. genetic, dietary and lifestyle.

While predicting longevity compounds from gene expression profiles using gene expression signatures proved to be successful in this study, this approach has additional limitations. First, the prediction scores from short-term treatment were not proportional to the effect size of the compounds on mouse lifespan. In fact, several compounds that had strong positive associations with biomarkers of lifespan-extending interventions and long-lived species after one month treatment in young mice failed to extend lifespan in old C57BL/6JN mice (e.g., GDC-0941 and ascorbyl-palmitate). As additional interventions are tested in mice, longevity biomarkers can be updated to improve prediction accuracy of the drugs. In addition, integration of other omics readouts and lifespan signatures from human tissues may further improve the predictive power of the developed approach. For example, protein phosphorylation of the MAPK pathway was predictive of longevity effects among compounds tested by ITP^23^, long non-coding RNA signatures predicted many mTOR and PI3K inhibitors that we evaluated here^24^, and integrating human brain aging signatures and CMAP predictions identified a few compounds that extended lifespan in our study^25^.

Another technical limitation is the number of compounds for which gene expression patterns are available. Although the number of compounds in the CMAP database is impressive (5,157 at the time of this study), it is far from what would be desirable. For example, the number of drugs that reach clinical stage is over 7,934 according to Drug Repurposing Hub (repo-hub.broadinstitute.org/repurposing), and drug screening libraries can include over 50,000 compounds (e.g. https://med.stanford.edu/htbc/compounds.html). Also, compounds identified in our study preferentially extended the lifespan of males. This is similar to the findings from ITP, where nine out of thirteen successful compounds extended male lifespan only^26,27,28,29^, and one compound was efficient in females only^30^. Short-term treatments used to identify most promising molecules in our pipeline were focused on male mice, which might have biased our compound selection. Indeed, compounds may have sex-specific effects on tissue gene expression profile, as demonstrated by a distinct effect of GDC-0941 on liver gene expression in male and female UM-HET3 mice (Fig. 3C). Including both sexes in screening might be beneficial for the identification of longevity compounds that work in both sexes.

We additionally would like to note that it is unknown which omics signatures are most informative. We relied on gene expression patterns in this study, whereas some groups prefer protein-based patterns. There is generally a lower than expected correlation between gene expression and protein levels in cells - across-gene correlations across multiple species demonstrated *r* values ranging from 0.39 to 0.79 ^31^. Perhaps, multi-omic approaches would be most useful in the future. Finally, we would like to stress that drug-induced changes to the transcriptome are only loosely predictive of drug effects in other cell types and most importantly in mice. Our work suggests that there is a low value in using CMAP gene expression data to identify drugs that extend lifespan in mice. It would be greatly beneficial for the biomedical community to develop a dataset of mice subjected to numerous interventions (drugs, diets, lifestyles) characterized by multi-omics approaches, including proteomics, transcriptomics, metabolomics, phosphophoproteomics, ATAC-seq and DNA methylation profiles from tissues with low-overlapping functions, especially these tissues that are safest to sample in future clinical trials (e.g. muscle, immune cells, reproductive tissues, skin). Such a dataset may be exceptionally useful, as opposed to cell-based datasets like CMAP, for the discovery of factors that influence complex phenotypes such as lifespan and incidence of disease.

Overall, we describe a platform for the identification of lifespan-extending interventions based on gene expression signatures of longevity and short-term tests in young mice that yielded 5 new compounds with significant lifespan-extending effects in mice, including vorinostat, selumetinib, LY294002, celastrol, and AZD8055. The identification of several new longevity interventions increases the pool of compounds that may be tested in humans. For example, selumetinib is an FDA-approved cancer drug, decreasing the barrier for its tests in clinical trials. Notably, we used a lower concentration of this compound in mice than used in cancer treatment, a situation resembling the use of rapamycin. Our platform may be used for rapid discovery of additional interventions, not limited to chemicals, and the newly discovered interventions may be used for defining general principles of lifespan control and for potential tests in the clinical setting.

## Methods

### EXPERIMENTAL MODEL AND STUDY PARTICIPANT DETAILS

#### Cell culture of primary hepatocytes

PHHs were commercially obtained from the MGH Cell Resource Core. The cells were cultured in a modified DMEM-based medium (Gibco, 11965092) optimized for hepatocyte maintenance. Prior to cell seeding, 96- and 384-well plates were coated with collagen by applying a 1:50 dilution of 1.25 mg/mL collagen in PBS, followed by a one-hour incubation at 37°C. After rinsing off excess collagen with PBS, the plates were air-dried. Hepatocytes were then seeded and allowed to attach for 24 hours before treating with compounds. Each compound was dissolved in DMSO and applied to PHHs at final concentrations of 1.11 µM, 3.33 µM, and 10 µM for 24 hours. Following the treatments, cells were either assayed for viability using the CellTiter-Glo detection reagent or lysed with TCL Lysis Buffer (Qiagen) for L1000 analysis ^32^.

#### Animals

##### UM-HET3 short-term treatment

3-month old UM-HET3 mice were bred at University of Michigan Medical School and subjected to diets containing compounds predicted via Connectivity Map (CMap): AS-703026 (30 ppm, as in ^33^, ascorbyl-palmitate (6.3 ppm, as in ^34^), AZD8055 (20 ppm, as in ^35^), celastrol (8 ppm, as in^36^), colforsin (forskolin) (8.3 ppm, as in ^37^), enzastaurin (30 ppm, as in ^38^), erismodegib (20 ppm, as in ^39^), everolimus (5 ppm, as in ^40^), GDC-0941 (30 ppm, as in ^41^), huperzine-a (0.2 ppm, as in^42^), kinetin-riboside (20 ppm, as in ^43^), KU-0063794 (10 ppm, as in ^44^), linifanib (10 ppm, as in ^45^), LY-294002 (100 ppm, as in ^46^), myricetin (150 ppm, as in ^47^), nifedipine (10 ppm, as in ^48)^, NVP-BEZ235 (50 ppm, as in ^49^), pyrvinium-pamoate (50 ppm, as in ^50^), rilmenidine (10 ppm, as in 51), selumetinib (50 ppm, as in ^52^), tert-butylhydroquinone (833.3 ppm, as in ^53^), trichostatin-a (1 ppm, as in ^54^), valdecoxib (5 ppm, as in ^55^), vorinostat (167 ppm, as in ^56^), and wortmannin (1.5 ppm, as in ^57^) for 1 month. Liver and kidney samples were taken from treated mice along with their untreated sex- and age-matched littermates, which were fed ad libitum.

##### Lifespan and healthspan assays in UM-HET3 mice

To prepare UM-HET3 mice used in the study, we mated young CByB6F1/J females and C3D2F1/J males (Jackson Laboratory), which were in turn obtained by mating individual inbred strains. UM-HET3 pups were weaned at 20-25 days of age and fed 5053 diet (TestDiet) *ad libitum* until 6 months of age. There were a maximum of 5 mice per cage. Mice were assigned to treatment groups so that the mean size of the litter they came from was similar across groups, because litter size has an effect on mouse lifespan^58^. Subsequently, mice were subjected to custom diets containing drugs (TestDiet) using 5053 as the base or the 5053 diet as control and followed until death or moribund state. Some mice were used in various healthspan assays and euthanized at 15 or 29 months of age; these mice were excluded from the survival curve.

##### Lifespan analysis using C57BL6/JN mice

24-25-month-old C57BL/6JN mice (both sexes) were obtained from the Aged Rodent Colony of the National Institute on Aging. After a few weeks of acclimation, animals were assigned to treatment groups based on frailty index score and body weight, so that the baseline mean and deviation of these parameters were similar across groups. In each group, 20-24 mice were subjected to custom diets containing drugs (TestDiet) using 5053 as the base or the 5053 diet as control and followed until death or moribund state. Our control group included 44 animals.

Drugs were purchased from Adooq or Medchem. All diets were irradiated after the drugs were incorporated into the chow. Mice that died as a result of fighting were excluded from survival analyses. Diets were replaced in the cages every two weeks. Mice were monitored daily until their natural death or euthanasia due to moribund state. Criteria for moribund state was labored breathing, reluctance to move, body condition score of 2 or lower, distended abdomen in combination with any of the criteria mentioned above, tumors over 2 cm, or eye tumors. Animals for cross-sectional studies were euthanized with CO_2_ followed by cervical dislocation. All experiments using mice were performed in accordance with institutional guidelines for the use of laboratory animals and were approved by the Brigham and Women’s Hospital and Harvard Medical School Institutional Animal Care and Use Committees (IACUC).

### METHOD DETAILS

#### RNA sequencing

Total RNA was extracted from snap-frozen livers using the Direct-zol RNA Miniprep Kit (Zimo) following manufacturer’s instructions. RNA was eluted with 50 ul of RNAse-free water. RNA concentration was measured with Qubit using an RNA HS Assay kit. Libraries were prepared with TruSeq Stranded mRNA LT Sample Prep Kit according to TruSeq Stranded mRNA Sample Preparation Guide, Part # 15031047 Rev. E. Libraries were quantified using the Bioanalyzer (Agilent) and sequenced with Illumina NovaSeq6000 S4 (2×150bp) (reads trimmed to 2×100bp) to obtain 20 million read depth coverage per sample. The BCL (binary base calls) files were converted into FASTQ using the Illumina package bcl2fastq. Fastq files were mapped to the mm10 (GRCm38.p6) mouse genome, and gene counts were obtained with STAR v2.7.2b^59^.

#### Frailty index

Frailty index was measured as described in the original paper^60^; frailty index score was calculated per the original protocol. Briefly, 31 parameters were visually assessed and graded as 0, 0.5 or 1, with the exception for weight and temperature that were quantitatively measured with scale and thermometer. Quantitative variables for weight and temperature were then converted into 0, 0.5 and 1 based on their deviation from means in the untreated group. Frailty index score was calculated as a mean value across all 31 parameters. C57BL/6JN mice were evaluated by two operators, and parameters that were significantly different between them were excluded for all mice from the analysis (breathing rate, gate disorders, piloreaction), although including all parameters did not significantly alter our results. The same cages were assigned to the same researcher in longitudinal assessment to ensure reproducibility. Researchers were not blinded to the mouse groups, but the results were collected before it was clear if drugs extended lifespan and the researcher had no prior hypothesis of which of these compounds will be effective (if any).

#### Gait speed

Mice were positioned in the regular mouse cages on the opposite side of a food container. Recording started with the SprinTimer app (App Store), and upon the ‘‘GO” signal from the app, mice were let run voluntarily to the other end of the cage. In the vast majority of cases, mice run towards the food container, presumably to hide. The finish line was marked by the colored tape at the end of the cage. When mice reached the finish line, recording was stopped and the time to finish was quantified based on the pictures from the app. The time of finish was determined as the time when the mouse nose crossed the finish line. Each mouse was let run four times and the median value was calculated. An attempt was considered successful if the mouse crossed the finish line without standing or turning 90 degrees or more during the run. Fifteen seconds were recorded If the mice failed to finish the run within 15 seconds.

#### Glucose and insulin tolerance tests

For the glucose tolerance test, mice were placed into food-free cages for 16 hours (usually from 7 pm until 11 am next day). The next day, mice were weighed, marked, bled from the tail and the fasting blood glucose level was measured with a glucometer. Filtered 10% glucose in sterile saline solution (Sigma) was injected i.p. at a final dose of 1 mg/kg of body weight. Injections were done at 30-second intervals between the mice. Glucose level was measured again at 20, 40, 60 and 120 minutes after the injection following the same 30-second interval between the mice. Insulin tolerance test was done similarly, with the difference that mice fasted for 6 hours (usually from 10 am to 4 pm), and were injected with 0.75 U/kg insulin (Eli Lilly) in sterile saline solution (Sigma).

#### Grip strength

All mice were brought to the experimental room at least 30 min prior to testing for acclimation. The grip was measured for fore limbs using a grid with Grip Strength Meter (Bioseb, Model GT3). Mice were gently lowered over the top of the grid so that front paws could grip the grid. Mice were allowed to attach to the grid properly before pulling it away. The torso was kept parallel to the grid, and the mouse was pulled back gently and horizontally until the grip was released down the complete length of the grid. Only pulls in which mice showed resistance to the experimenter were taken into account. Four measures were taken. The grid was cleaned with 70% ethanol between mice.

#### Rotarod

Mice were brought to the experimental room at least 30 min prior to testing (for acclimation). Animals were first exposed to the apparatus (Stoelting; Ugo Basile Apparatus) in a 5-minute habituation training session at a constant speed of 4 rpm. When a mouse fell or turned 180 degrees on the rod during this training session, it was repositioned correctly on the apparatus until the mouse completed the 5 min session. After all mice were tested during the habituation session, mice were tested in the accelerating test. The time between the start of the habituation and the test session was 2 hours. During this test session the speed was set to increase steadily at 4 rpm within a 5 min period. Mice were placed on the rod at the 4 rpm speed. When mice reversed the position or grabbed the roller and rotated with the rod, they were gently repositioned. Once the mouse fell off the rotarod, the timer was automatically stopped.

#### Open field

Mice were brought to the experimental room at least 30 min prior to testing for acclimation. Then, mice were placed in the test arena for an 1-hour session. The test consisted of sessions which were automatically started by placing animals individually in the center of the arena. Animals were allowed free exploration of the environment for 1 hour with recordings in 5 min bins. During the session, animals did not have access to food or water. A computer-assisted infra-red tracking system was used to record the number of beam breaks (activity) and user-defined zone entries. The test arena (Med Associates, St. Albans, VT, ENV-510) was made of Plexiglas and consisted of a square 27.3 cm x 27.3 cm base with 20.3 cm high walls. All the walls were clear. Light was kept dim. A power station-Med Associates SG506 was connected to all ENV-510 boxes. Between sessions all feces were removed and the chambers were thoroughly cleaned with 70% ethanol and water.

#### Comprehensive laboratory animal monitoring system (CLAMS™)

Mice were brought to the experimental room at least 30 min prior to testing for acclimation. Twenty-five-month-old treated and control mice were individually placed in CLAMS™ cages and monitored over a 4-day period, with animals being checked daily. If a mouse refused to eat or drink for 1 day, it was removed from the experiment. Food and water consumption were measured directly as accumulated data. The hourly file displayed all measurements for each parameter: VO2 (volume of oxygen consumed, mL/Kg/hr), VCO2 (volume of carbon dioxide produced, mL/Kg/hr), RER (respiratory exchange ratio), heat (Kcal/hr), accumulated food (grams), accumulated drink (grams), XY total activity (all horizontal beam breaks in counts), XY ambulatory activity (minimum 3 different, consecutive horizontal beam breaks in counts) and Z activity (all vertical beam breaks in counts). The data were recorded during the 60-second sampling period. Data was exported using Oxymax software as .csv files and analyzed in Rstudio.

#### Necropsy analysis

Mice were euthanized with CO_2_ followed by cervical dislocation. The chest and abdomen were opened, and the body was immersed into formalin solution and stored at 4°C until further analysis. For necropsy analysis all organs, including small endocrine organs, were dissected, trimmed at 5 mm thickness and embedded in paraffin blocks. Paraffin blocks were sectioned at 5 μm and stained with hematoxylin and eosin. The slides were examined blindly by a pathologist at the Harvard Rodent Histopathology Core.

#### Flow cytometry

mAbs used for staining included: anti-CD19 [6D5], anti-CD11b [M1/70], anti-CD3 [17A2], anti-CD23 [B3B4], anti-CD21 [7E9], anti-CD24 [M1/69], anti-CD29 [HMβ1-1], and anti-CD45 [30-F11] (all from Biolegend). Dead cells were excluded by DAPI staining. Data was collected on a Cytek DXP11 and analyzed by FlowJo software (BD). Spleens were gently pressed over 40 um cell strainer snap cap (Corning) to get single-cell suspensions. Cell suspensions were centrifuged at 4°C, 300 g for 5 min, resuspended in 1 ml of red blood lysis buffer for 10 min on ice, washed once with 1 ml of FACS buffer (1% FBS in PBS), stained in 100 ul of antibody solution (2 ng/ul of each antibody) at 4°C for 20 min protected from light, washed again, resuspended in 1 ml of FACS buffer, filtered into tubes with cell strainer snap cap (Corning), and analyzed with flow cytometry. One hundred thousand events were recorded.

### QUANTIFICATION AND STATISTICAL ANALYSIS

#### in silico screening with CMAP

*in silico* screening was performed with the Clue platform from CMAP. For each of the 9 gene expression signatures of longevity (5 signatures of lifespan-extending interventions^13^ and 4 signatures of long-lived mammals^14^), we selected top 150 differentially expressed genes (Benjamini-Hochberg adjusted p-value < 0.05) with the highest absolute size of effect (|logFC|), and provided lists of up- and downregulated genes as query inputs to the Clue platform. To adjust for possible variation in the gene expression response across cell lines, for every compound we pooled predicted association scores across all available cell types and concentrations. For each query, we standardized normalized enrichment scores (NES) to ensure comparability across signatures. Consistency of NES across signatures was assessed with pairwise Spearman correlation. To integrate results across intervention biomarkers and species, we computed aggregated longevity scores, defined as the mean of standardized NES derived from signatures of maximum lifespan in mice (“Max lifespan”) and long-lived species across organs (“Pooled ML”).

#### Gene expression signatures of compounds in PHH

Gene expression profiles of PHH were obtained using the L1000 platform. Gene probes were mapped to Entrez ID, resulting in 13,433 detected genes. Alignment of transcriptomic profiles was assessed through density distributions. The presence of outliers was evaluated using PCA. The consistency of compound-induced gene expression changes was assessed with Spearman correlation. Pairwise correlation coefficients were computed separately for sample pairs corresponding to the same compound and those corresponding to different compounds. The median correlation coefficient between groups was compared with the Wilcoxon rank sum test.

Differentially expressed genes for each compound and dosage were identified with limma^61^. DMSO-treated and untreated samples served as controls, with plate included as a covariate in the statistical model. The False Discovery Rate (FDR) correction based on Benjamini-Hochberg (BH) approach was performed to adjust for multiple hypothesis testing^62^. Genes with an adjusted p-value < 0.05 were considered statistically significant. Associations between the number of differentially expressed genes across treatments and dose or change in viability (log-ratio) were assessed using the Wilcoxon signed-rank test and Spearman correlation, respectively. To derive a gene expression signature of PHH toxicity, we applied a limma regression model with log-viability as the outcome value. Genes with an adjusted p-value < 0.05 were considered statistically significant. For GSEA, genes were ranked based on their −log10(p-value) multiplied by sign(logFC). HALLMARK, KEGG, and REACTOME ontologies from the Molecular Signature Database (MSigDB) were used as gene sets for GSEA. The GSEA was performed via the fgsea package in R with a multilevel splitting Monte Carlo approach and 5000 permutations. Calculated p-values were adjusted with BH approach. Pathways with BH-adjusted p-values < 0.05 were considered statistically significant.

To generate an aggregated compound signature in PHH across doses, we first excluded toxic doses, defined as those with log-viability below the lower bound of the 99.98% prediction interval for DMSO-treated and untreated cells. The remaining samples for each compound were pooled, and an aggregated gene expression signature was computed using the limma model as described above. After filtering out toxic doses, we obtained aggregated transcriptomic signatures for 99 compounds, which were used as inputs for signature association analysis. To assess the association of aggregated compound signatures in PHH with nine established signatures of lifespan-extending interventions and long-lived species, we employed a previously developed GSEA-based algorithm^13,14^. For each longevity signature, we selected the top 1,000 statistically significant genes (BH-adjusted p-value < 0.05) with the highest absolute effect size (|logFC| or |slope|). The NES obtained from this analysis were standardized, and aggregated NES was calculated in the same manner as for the CMAP screening.

#### RNA-seq analysis of tissues from young UM-HET3 after short-term treatment

RNA-seq data from liver and kidney samples were generated in two rounds. Raw expression data from each round and tissue were filtered separately, and genes with at least 5 reads in 50% of samples were kept for subsequent analysis. Gene annotations were mapped to Entrez IDs, and the resulting expression profiles were normalized using RLE normalization^63^. Differentially expressed genes for each compound and tissue were identified using custom models in R with edgeR^64^. To evaluate the pooled effect of each compound across tissues, gene expression data were pooled across tissues, and tissue type was included as a covariate in the statistical model. Multiple hypothesis testing was corrected using the BH method, with genes considered statistically significant at a BH-adjusted p-value < 0.05.

Functional GSEA was conducted for the pooled signature of each compound across tissues, following the approach described above. To identify toxicity-associated terms, an unpaired t-test was performed separately for each term, comparing the NES of toxic compounds (AS-703026 and wortmannin) against those of all other compounds. The resulting p-values were adjusted using the BH method, and pathways with BH-adjusted p-value < 0.05 were considered statistically significant.

To evaluate the consistency of gene expression changes induced by the same compound across liver, kidney, and PHH, we conducted the following pairwise analyses. For each pair of models (e.g., liver vs. kidney), we computed Pearson’s correlation between the logFC values induced by each compound in the first model and those in the second model. Next, for each compound in the first model, we ranked all compounds in the second model based on their correlation coefficient and recorded the rank of the matching compound. This process was repeated for every compound in the first model. To quantify consistency, for every rank *i*, we calculated the percentage of correctly matched compounds appearing within the top *i* hits from the second model and visualized it on the plot (Fig. 2B, Fig. S2A,B). Finally, we computed the area under the curve (AUC) and performed a permutation test to assess the statistical significance of the observed consistency between model pairs.

NES for individual longevity signatures and the aggregated longevity score were computed for gene expression changes induced by each compound in liver, kidney, and the pooled cross-tissue signature, as described above. The consistency of NES across individual longevity signatures and the consistency of aggregated longevity scores across models (tissues, PHH, CMAP) for matched compounds were evaluated using Spearman correlation. To derive the aggregated longevity score for each compound across all tested models, we first standardized longevity scores within each model. We then calculated a weighted mean of the normalized scores across models, assigning weights of 1 to both the CMAP and PHH models, and a weight √2 of to the murine tissue data. This weighting scheme was selected to reflect both the greater biological relevance of the tissue data and the use of two distinct tissues (liver and kidney) in generating the tissue-level scores.

#### RNA-seq analysis of tissues from UM-HET3 after long-term treatment

Raw expression data from liver samples of male and female UM-HET3 mice following long-term treatment were filtered, retaining genes with at least 10 reads in 25% of samples. Gene annotations were mapped to Entrez IDs and normalized using the RLE method. Data quality was evaluated via density distributions and PCA, leading to the exclusion of two outliers.

Genes differentially expressed in at least one of the treatment groups, were identified using ANOVA, incorporating sex and diet as covariates. P-values were adjusted with the BH method, and genes with BH-adjusted p-values < 0.05 were considered statistically significant. To assess organization of treatments according to their transcriptomic profiles, we performed hierarchical clustering of individual samples using Spearman correlation distance and complete linkage on ANOVA-identified genes (Fig. 5B). Genes differentially expressed in response to each compound were identified separately for males and females using edgeR. In males, genes associated with compound effects on expected maximum lifespan were identified via a regression model, with the 90th percentile of maximum lifespan (P90) as the outcome. Genes with BH-adjusted p-values < 0.05 were considered statistically significant. To evaluate consistency of compound-induced gene expression changes across sexes (Fig. 4C) and across treatment durations in males (1-month vs. 9-month treatment) (Fig. S4D), we calculated Spearman correlation using the union of the top 500 genes with the lowest p-values for each comparison.

Functional GSEA was conducted for individual compound signatures (separately for males and females), for signature of maximum lifespan in UM-HET3 males, for established liver signatures of lifespan-extending interventions and long-lived species, and for established liver and multi-tissue aging signatures in mice and rats^14^, as described previously. Hierarchical clustering of enriched functions for visualization (Fig. 5F, Fig. S4C) was based on NES, using complete linkage and Manhattan distance. Functions (i) significantly enriched (adjusted p-value < 0.05) in multiple signatures and (ii) representing distinct biological processes, as determined manually and via gene set overlap, were selected for visualization. The whole list of statistically significant enriched functions is available in Table S3.

#### DNA methylation analysis

We analyzed DNA methylation using several epigenetic clocks based on the mammalian methylation array (HorvathMammalMethylChip40)^65^. The array and the related clocks have been employed in numerous previous studies including by the Mammalian Methylation Consortium^66,67^. This array is commercially available through the Epigenetic Clock Development Foundation (Torrance, USA). All R software scripts and clock algorithms have been previously published and are accessible via various publications and the MammalMethylClock R package^68^.To ensure the robustness of our findings, we simultaneously evaluated multiple epigenetic clocks. The consistent indication of epigenetic rejuvenation across different clocks suggests the reliability of our results, independent of specific clock construction methods. For instance, heart tissue was assessed using three clocks developed specifically for murine heart: an elastic net regression-based clock^69^, an “interventional” clock based on CpGs related to validated life-extending interventions^69^, and the LUC clock, which uses cytosines associated with lifespan^70^.

## Supporting information

List of diets tested in mice for a short term

NES generated using longevity signatures and applied across in vitro and in vivo studies

Pathway enrichments across in vitro and in vivo studies

Detected lesions in C57BL6/JN mice

Lifespan statistics for C57BL6/JN mice

Lifespan statistics for UM-HET3 mice

## Competing interests

The Regents of the University of California are the sole owner of patents and patent applications directed at epigenetic biomarkers and the mammalian methylation array platform for which Steve Horvath is a named inventor; SH is a founder and paid consultant of the non-profit Epigenetic Clock Development Foundation that licenses these patents. SH is a Principal Investigator at the Altos Labs, Cambridge Institute of Science, a biomedical company that works on rejuvenation. AVS and AT were employed by Retro Biosciences after completion of experiments and analysis of data and own shares of the company. AVS, AT and VNG have applied for a patent application related tot his work.

## Acknowledgements

Supported by grants from the National Institute on Aging.

---

**Figure S1.**
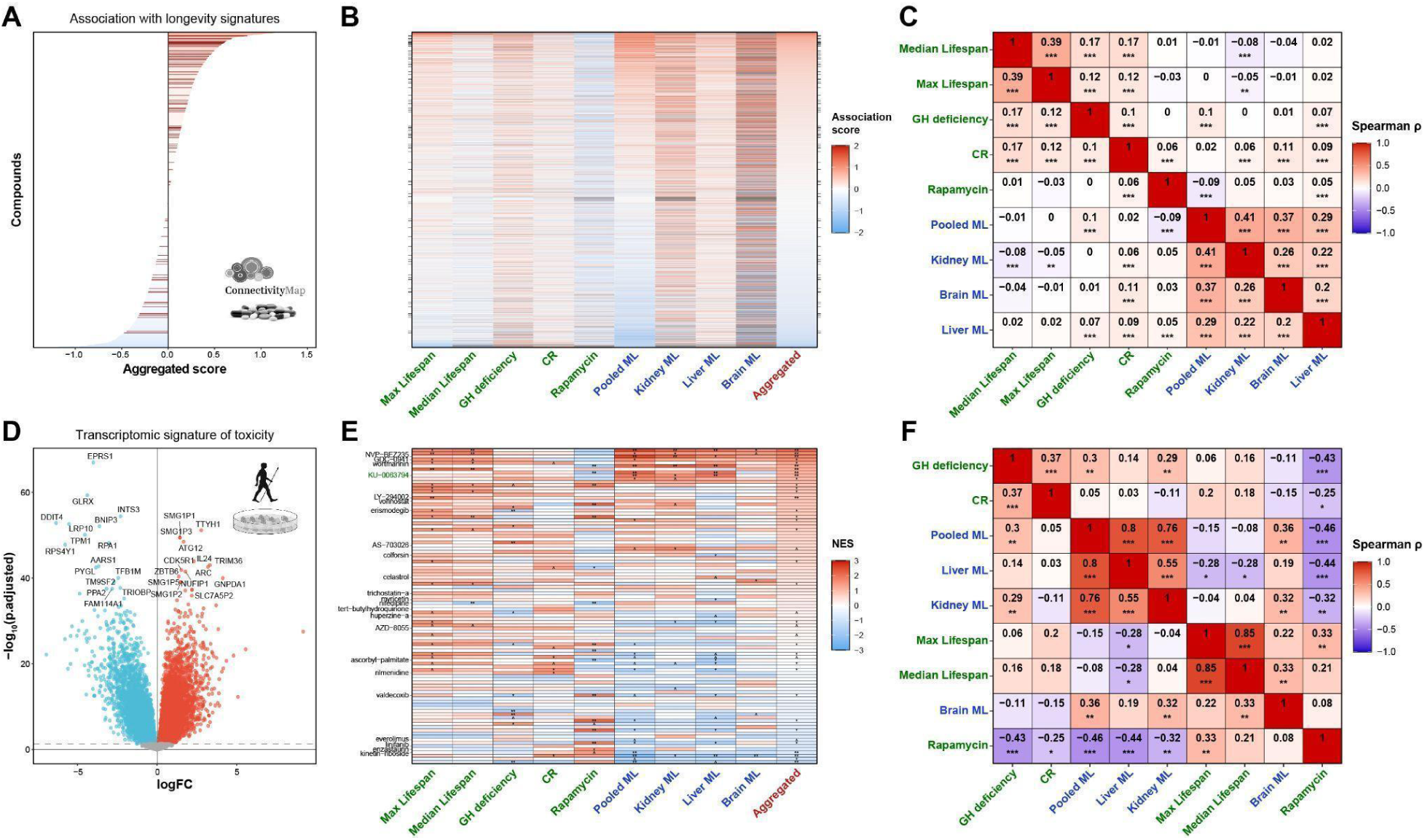
related to Figure 1. Associations of gene expression changes induced by compounds in human cells with signatures of longevity. **(A) Aggregated longevity association score of chemical compounds according to the Connectivity Map (CMAP) database.** Compounds are sorted based on the average association score of their gene expression profiles across various human cell lines with signatures of maximum lifespan in mice and long-lived species across organs. Compounds selected for screening in primary human hepatocytes (PHH) are shown with red lines. **(B) Associations of gene expression profiles of chemical compounds in CMAP with individual signatures of lifespan-extending interventions in mice (green) and long-lived mammalian species (blue).** Every column corresponds to the individual signature. Aggregated association score (presented on Fig. S1A) is shown in red and is calculated as the mean of association scores for signature of maximum lifespan in mice (“Max Lifespan”) and signature of long-lived species across organs (“Pooled ML”). Compounds selected for screening in PHH are shown with ticks on sides of the heatmap. The output of the association test is in Table S2. **(C) Spearman correlation between association scores of compounds in CMAP calculated based on various individual signatures of longevity.** Signatures of lifespan-extending interventions and long-lived species are labeled in green and blue, respectively. Correlation coefficients and corresponding BH-adjusted p-values are shown with text and asterisks, respectively. **(D) Volcano plot of genes associated with toxicity (cell death) in PHH.** Top 30 genes with the lowest BH-adjusted p-value are labeled with gene symbols. FDR threshold of 0.05 is shown with dotted line, significant genes (BH-adjusted p-value < 0.05) positively and negatively associated with cell death are colored in red and blue, respectively. **(E) Associations of gene expression profiles of chemical compounds in PHH with individual signatures of lifespan-extending interventions in mice (green) and long-lived mammalian species (blue).** Every column corresponds to the individual signature. Aggregated association score (presented on Fig. 1G) is shown in red. Compounds selected for screening in murine tissues are labeled in text. KU-0063794, established to extend lifespan in old mice in our previous study, is labeled in green. BH-adjusted p-values are denoted by asterisks. The output of the association test is in Table S2. **(F) Spearman correlation between association scores of compounds in PHH assay calculated based on various individual signatures of longevity.** Signatures of lifespan-extending interventions and long-lived species are labeled in green and blue, respectively. Correlation coefficients and corresponding BH-adjusted p-values are shown with text and asterisks, respectively. NES: Normalized Enrichment Score; Max: Maximum; CR: Caloric restriction; GH: Growth hormone; ML: Maximum Lifespan.

**Figure S2.**
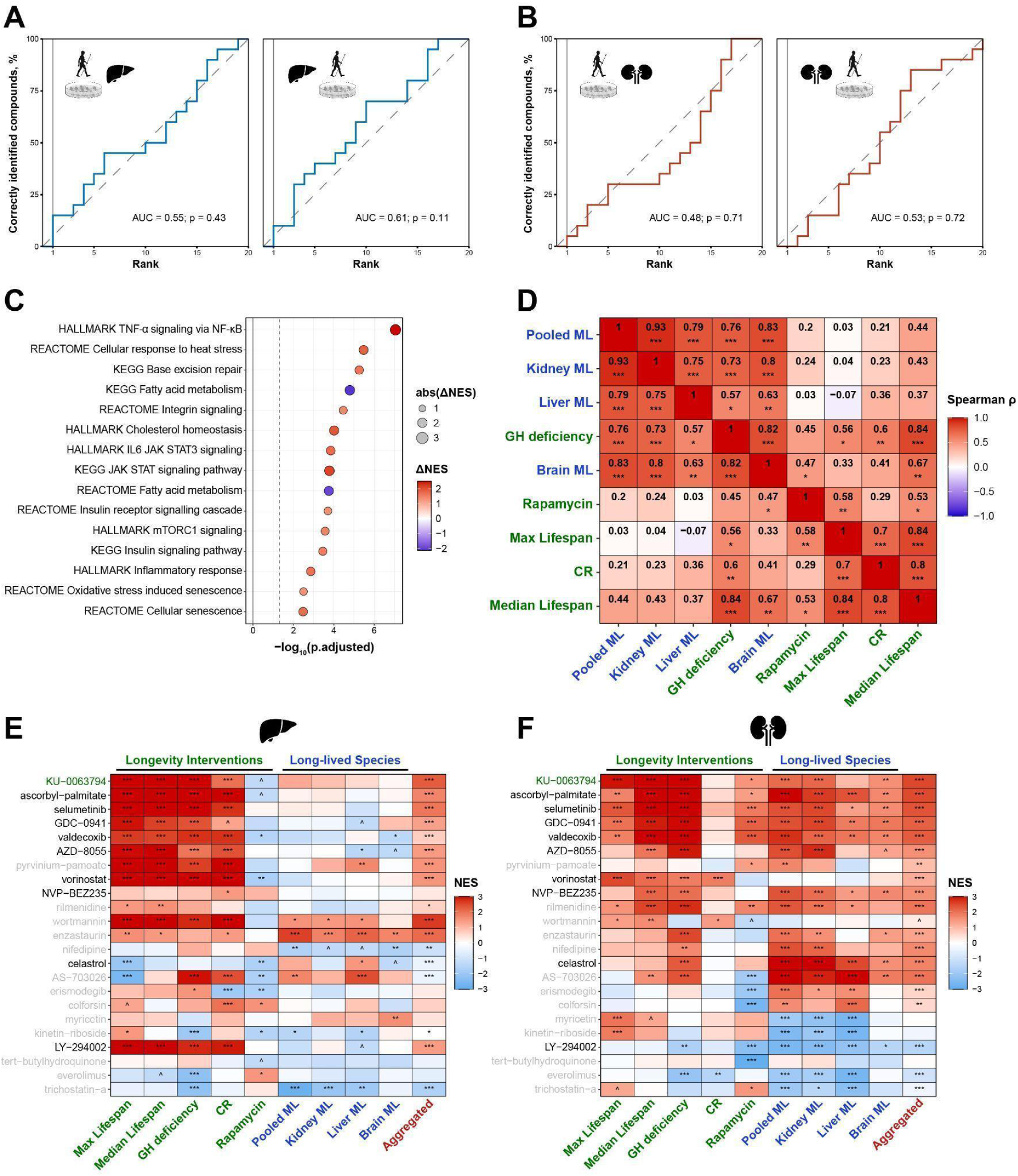
related to Figure 2. Association of gene expression changes induced by 1-month application of compounds in murine liver and kidney with signatures of longevity. **(A)-(B) Consistency between gene expression changes induced by individual compounds in primary human hepatocytes and mouse liver (A) or kidney (B).** The y axis reflects the percentage of compounds, whose gene expression profiles in mouse tissue (left) and PHH (right) are among the top correlated with the reference profiles of the same molecules in the other biological model. The x-axis indicates the number of top-ranked positions used to identify a match in the ranking list. Pearson correlation was used to rank the compounds. AUC and permutation test p-value are shown in text. **(C) Enriched pathways that distinguish gene expression profiles of toxic compounds (AS-703026 and wortmannin) from other treatments.** Difference in average NES between toxic and other compounds is shown with color and dot size. Pathways positively and negatively associated with toxic compounds compared to other treatments are colored in red and blue, respectively. Statistical significance of NES difference between the groups (BH-adjusted p-values) was assessed with unpaired t-test and plotted on x axis. The whole list of enriched functions is in Table S3. **(D) Spearman correlation between NES of compound gene expression profiles in murine tissues estimated for various individual signatures of longevity.** Association was assessed for pooled gene expression changes across liver and kidney. Signatures of longevity interventions and long-lived species are labeled in green and blue, respectively. Correlation coefficients and corresponding BH-adjusted p-values are shown with text and asterisks, respectively. **(E)-(F) Association of gene expression changes induced by tested compounds in liver (E) and kidney (F) with signatures of lifespan-extending interventions (green) and long-lived mammalian species (blue).** Cells are colored based on the NES calculated for gene expression profiles of compounds separately for each tissue. Aggregated longevity score is labeled in red. KU-0063794, established to extend lifespan in old mice in our previous study, is labeled in green, whereas compounds chosen for a survival study in old C57BL/6JN mice are labeled in black. BH-adjusted p-values are denoted by asterisks. The output of the association test is in Table S2. ^ p.adjusted < 0.1; * p.adjusted < 0.05; ** p.adjusted < 0.01; *** p.adjusted < 0.001. AUC: Area Under the Curve; Max: Maximum; CR: Caloric restriction; GH: Growth hormone; ML: Maximum Lifespan; NES: Normalized Enrichment Score.

**Figure S3.**
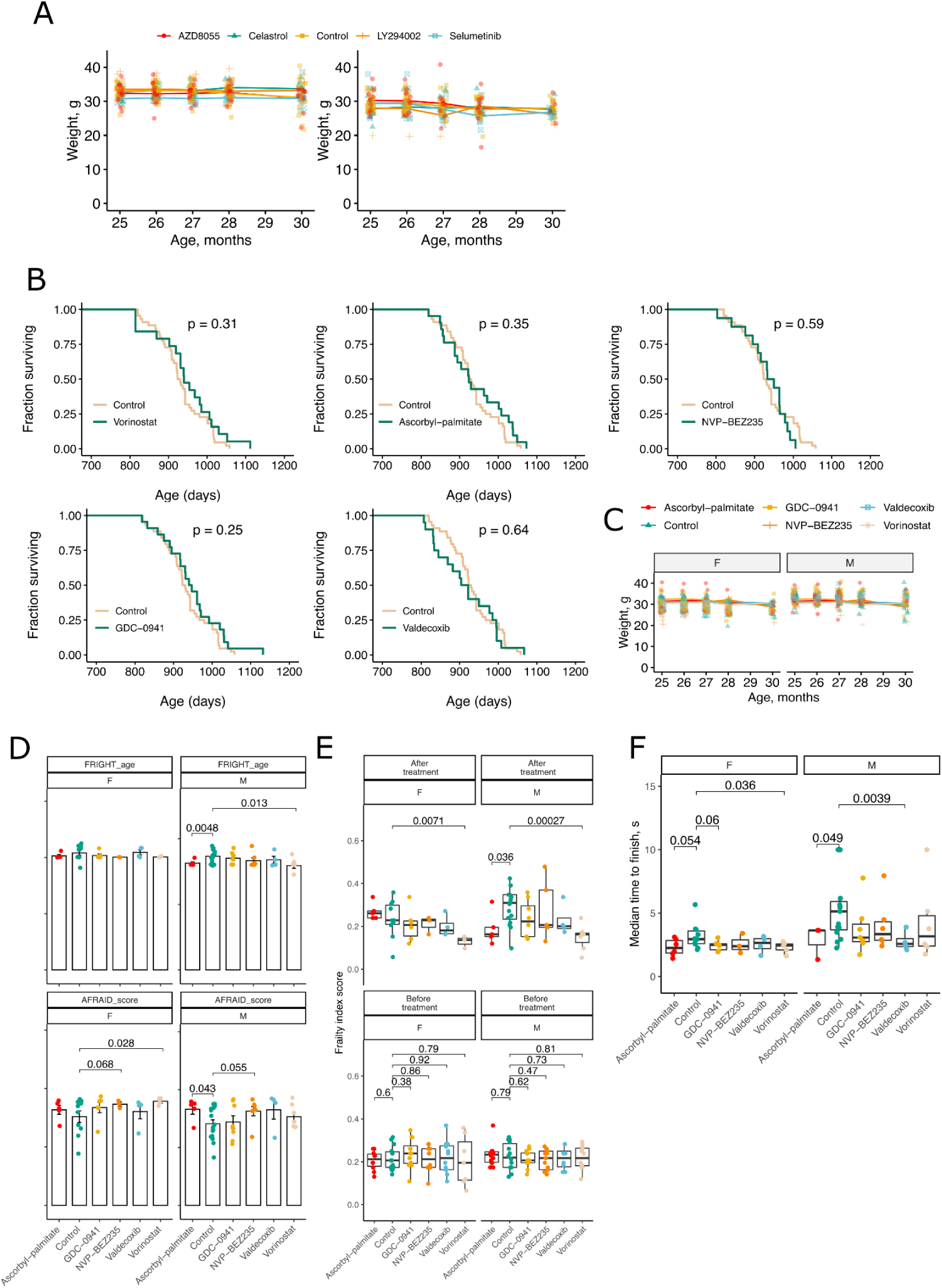
related to Figure 3. Lifespan and healthspan of aged C57BL/6JN mice subjected to diets containing predicted longevity interventions. **(A) Body weight of** C57BL/6JN **males and females treated with longevity candidate compounds in food.** Each dot is a mouse. **(B) Lifespan of** C57BL/6JN **mice treated with candidate longevity compounds in food at old age.** Survival of both sexes following treatment with 1000 ppm vorinostat, 38.5 ppm ascorbyl-palmitate, 50 ppm NVP-BEZ235, 50 ppm GDC-0941, or 5 ppm Valdecoxib, all compared to untreated control. P-values were calculated with a log-rank test. **(C) Body weight of** C57BL/6JN **males and females treated with candidate longevity compounds in food.** Each dot is a mouse. **(D) Frailty index clocks measured in 31 months old** C57BL/6JN **mice after 5 months of treatment.** FRIGHT age predicts age of the mice based on frailty index measures, and AFRAID score predicts survival time. P-values were calculated with a two-sided Student T-test. **(E) Frailty index score measured in 31 months old** C57BL/6JN **mice after 5 months of treatment.** Mice were subjected to diets from 24-25 months of age and the frailty score was measured before treatment and at 31 months of age in both sexes. P-values were calculated with a two-sided Student T-test. **(F) Median time to finish the gait speed test measured in 31-month-old** C57BL/6JN **mice after 5 months of treatment.** Gait speed was measured four times per mouse, and the median value was plotted and calculated. P-values were calculated with a two-sided Student T-test.

**Figure S4.**
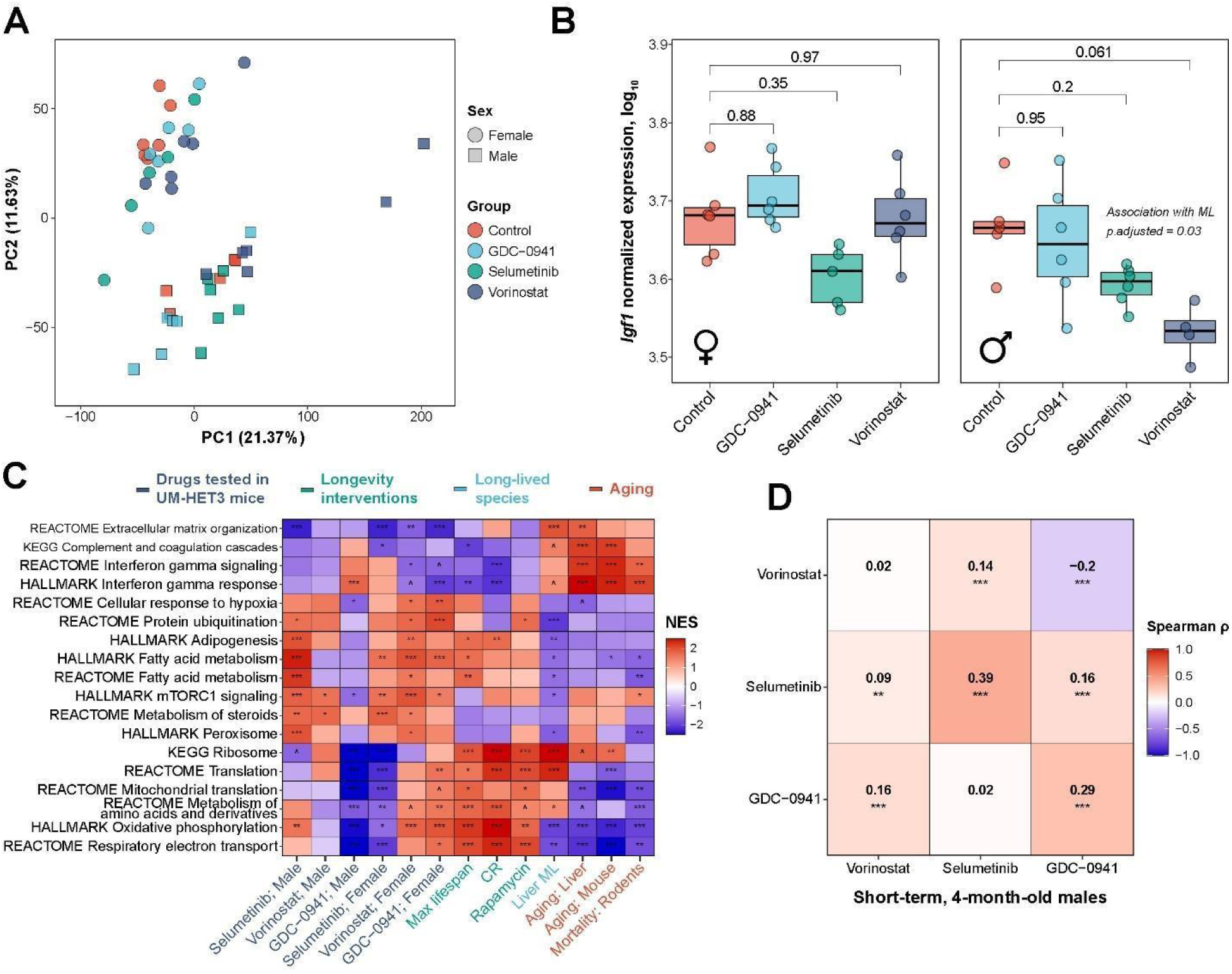
related to Figure 5. Gene expression response of UM-HET3 mice to long-term treatment with selected longevity-associated compounds. **(A) Principal component analysis of mouse liver samples.** Each dot represents an individual biological replicate. Females and males are shown in circles and squares, respectively. Color represents the treatment group. Percentage of variance explained by first and second principal components is provided in brackets. Two outlier samples with PC1 values greater than 100 were excluded from further analysis. n=5-6 per treatment group per sex. PC: Principal component. **(B) Normalized expression of *Igf1* in livers of age-matched female (left) and male (right) mice subjected to 9 months of oral treatment with selected compounds.** BH-adjusted p-values for pairwise comparisons between control and treated groups assessed with edgeR are labeled with numbers. Statistical significance of association between expression of the gene and maximum lifespan of male groups assessed with linear model is provided in text. **(C) Functional enrichment of gene expression changes associated with individual tested compounds in UM-HET3 male and female mice (indigo), along with established signatures of lifespan-extending interventions (green), long-lived species (lightblue), aging and mortality (red).** GSEA was performed using HALLMARKS, REACTOME, and KEGG ontologies. Only pathways statistically significantly enriched for a signature of maximum lifespan in UM-HET3 male mice (BH-adjusted p-value < 0.05) are shown. Normalized enrichment score (NES) is shown with color, and BH-adjusted p-values are denoted by asterisks. The whole list of enriched functions is in Table S3. **(D) Spearman correlation of gene expression changes induced by compounds in livers of UM-HET3 males after short-term 1-month (in columns) and long-term 9-month (in rows) treatments.** For every pairwise comparison, the union of top 500 genes with the lowest p-value was used to estimate a correlation coefficient. Pairwise Spearman’s ⍴ are shown in color and text, and BH-adjusted p-values are denoted with asterisks. Max: Maximum; CR: Caloric restriction; GH: growth hormone; ML: Maximum Lifespan. ^ p.adjusted < 0.1; * p.adjusted < 0.05; ** p.adjusted < 0.01; *** p.adjusted < 0.001. See also Figure S4 and Table S3.

**Figure S5.**
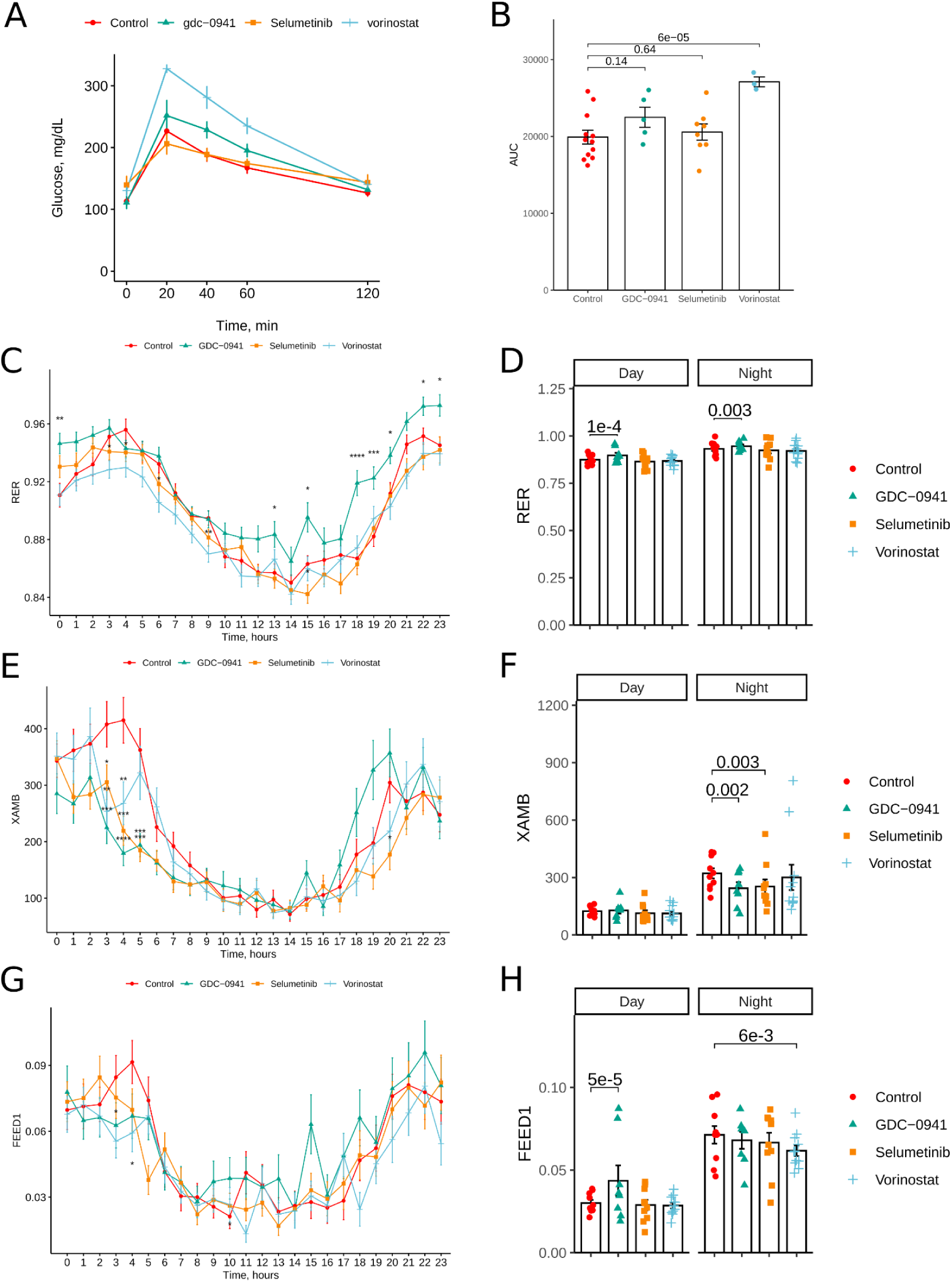
related to Figure 7. Measures of metabolic health in 6-months old and 22-25-months old UM-HET3 mice treated with longevity compounds starting at young age. **(A-B) Results of glucose tolerance test measured in 6-months old UM-HET3 mice.** Values are mean ± SD for both sexes combined. Barplots are areas under the curve (AUC) for the glucose tolerance test. P-values are ANCOVA adjusted for sex. Each dot is a mouse. **(C-D) Respiratory exchange ratio (RER) of 25-months old UM-HET3 female mice treated with longevity intervention and compared to untreated control.** Values are mean ± SD, averaged over a 24-72 hour period of recordings. Shadowed areas indicate dark time at the facility, when mice are active. Barplots are averaged RER values during day and night periods. P-values are calculated with repeated measures ANOVA. Each dot is a mouse. **(E-F) Horizontal ambulatory activity (XAMB) of 25-months old UM-HET3 female mice treated with longevity intervention and compared to untreated control.** Values are mean ± SD, averaged over a 24-72 hour period of recordings. Shadowed areas indicate dark time at the facility, when mice are active. Barplots are averaged XAMB values during day and night periods. P-values are calculated with repeated measures ANOVA. Each dot is a mouse. **(G-H) Eating activity in grams (FEED1) of 25-months old UM-HET3 female mice treated with longevity intervention and compared to untreated control.** Values are mean ± SD, averaged over a 24-72 hour period of recordings. Shadowed areas indicate dark time at the facility, when mice are active. Barplots are averaged FEED1 values during day and night periods. P-values are calculated with repeated measures ANOVA. Each dot is a mouse.

**Figure S6.**
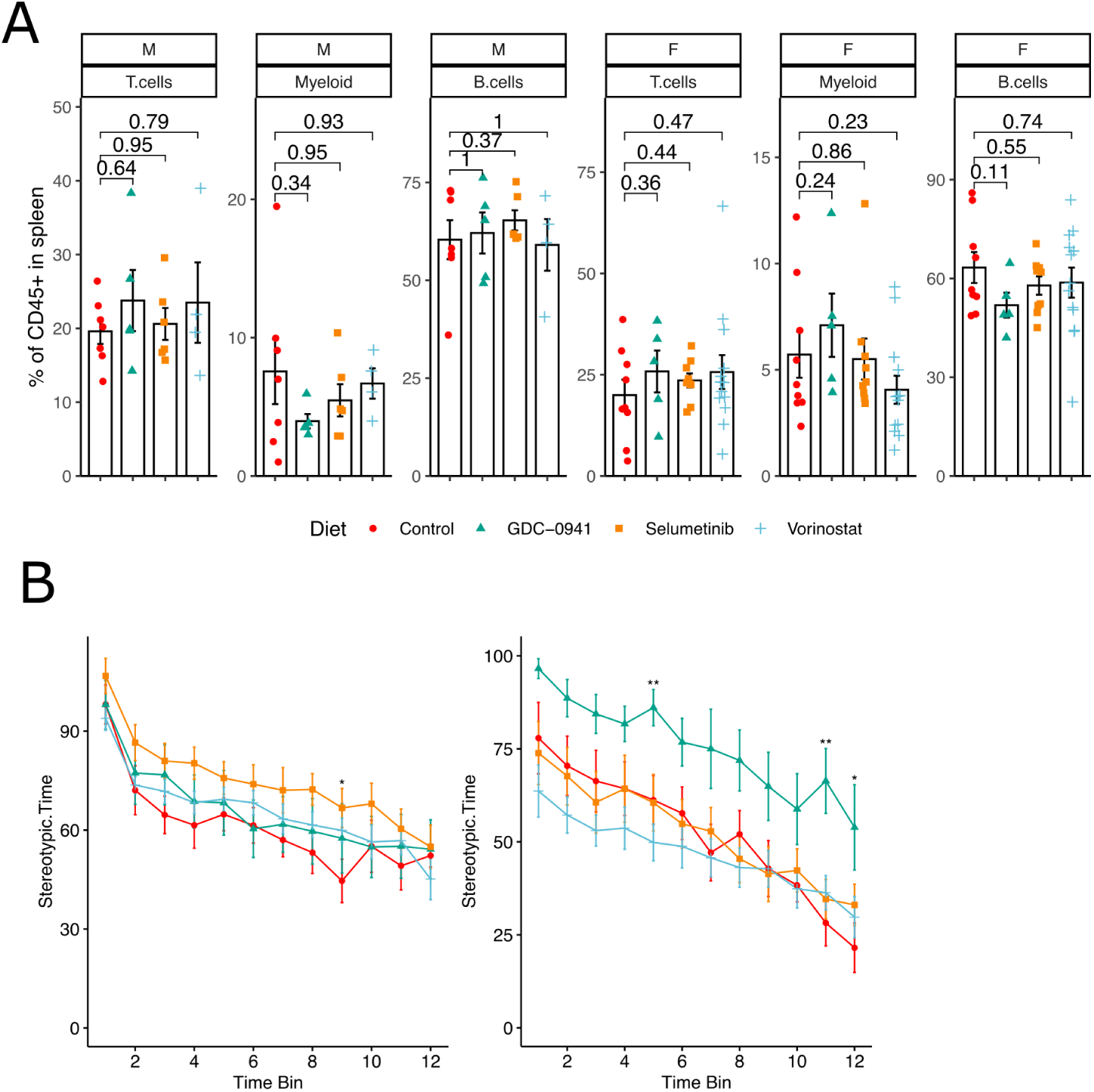
related to Figure 6. Measures of health in 22- or 28-month-old UM-HET3 mice treated with longevity compounds starting at 6 months of age. **(A) Proportion of B cells, T cells and myeloid cells measured in spleens of 28-month-old UMHET mice.** Females (F) and males (M) were treated with one of three longevity compounds and compared to untreated controls. Each dot is a mouse, dots are colored and shaped based on the treatment group. P-values were calculated with a two-sided Student T-test. **(B) Locomotor activity of 22-month-old UM-HET3 mice treated with longevity compounds and compared to untreated controls.** Mean values with standard deviation are shown for each treatment group in males and females separately. Each bin is a 5 min interval of a 1 hour recording session. Stereotypic time is the time mice spend crossing the arena during recordings.

